# Divergent disruption of brain networks following total and chronic sleep loss: a longitudinal fMRI study

**DOI:** 10.1101/2025.10.10.681651

**Authors:** Patrycja Scislewska, Arturo Cabrera Vazquez, Iwona Szatkowska, Halszka Kontrymowicz-Ogińska, Sophie Achard, Aleksandra Domagalik

**Author notes:** These authors contributed to this work equally (Sophie Achard, Aleksandra Domagalik).

## Abstract

**Study objectives:** Sleep loss significantly disrupts cognitive and emotional functioning, yet the neural consequences of different types of sleep deprivation remain unclear.

**Methods:** In a within-subject resting-state fMRI study, we examined how acute total sleep deprivation (TSD) and chronic sleep restriction (CSR) alter intrinsic functional brain organization in 28 healthy adults scanned under three conditions: rested wakefulness (RW), after one night of TSD, and after five nights of CSR.

To quantify network-level disruption, we applied graph-theoretical analyses, including a novel within-subject adaptation of the Hub Disruption Index and Covariate-Constrained Manifold Learning (CCML), an unsupervised embedding technique sensitive to subject-level covariates. Moreover, we assessed subjective sleep quality, sleepiness, and circadian traits.

**Results:** Both TSD and CSR were associated with a consistent reorganization of graph topology relative to RW. Furthermore, direct comparisons revealed that TSD and CSR affect different brain hubs. Regional changes in degree, closeness, and clustering coefficients were most prominent in subsystems of the default mode network, frontoparietal network, and cerebellum. These differences were also captured in CCML embeddings, supporting the hypothesis that acute and chronic sleep deprivation exert divergent effects on brain connectivity. Findings were robust across graph thresholds, brain atlases, and nodal metrics. Moreover, these results were further supported by the subjective measures - sleepiness was associated with reduced network integration in RW, and circadian phenotype emerged as a key determinant of individual sensitivity to sleep loss.

**Conclusions:** Our results show that TSD and CSR induce distinct alterations in brain functional organization, offering new insights into their neural impact.

**Graphical abstract:** 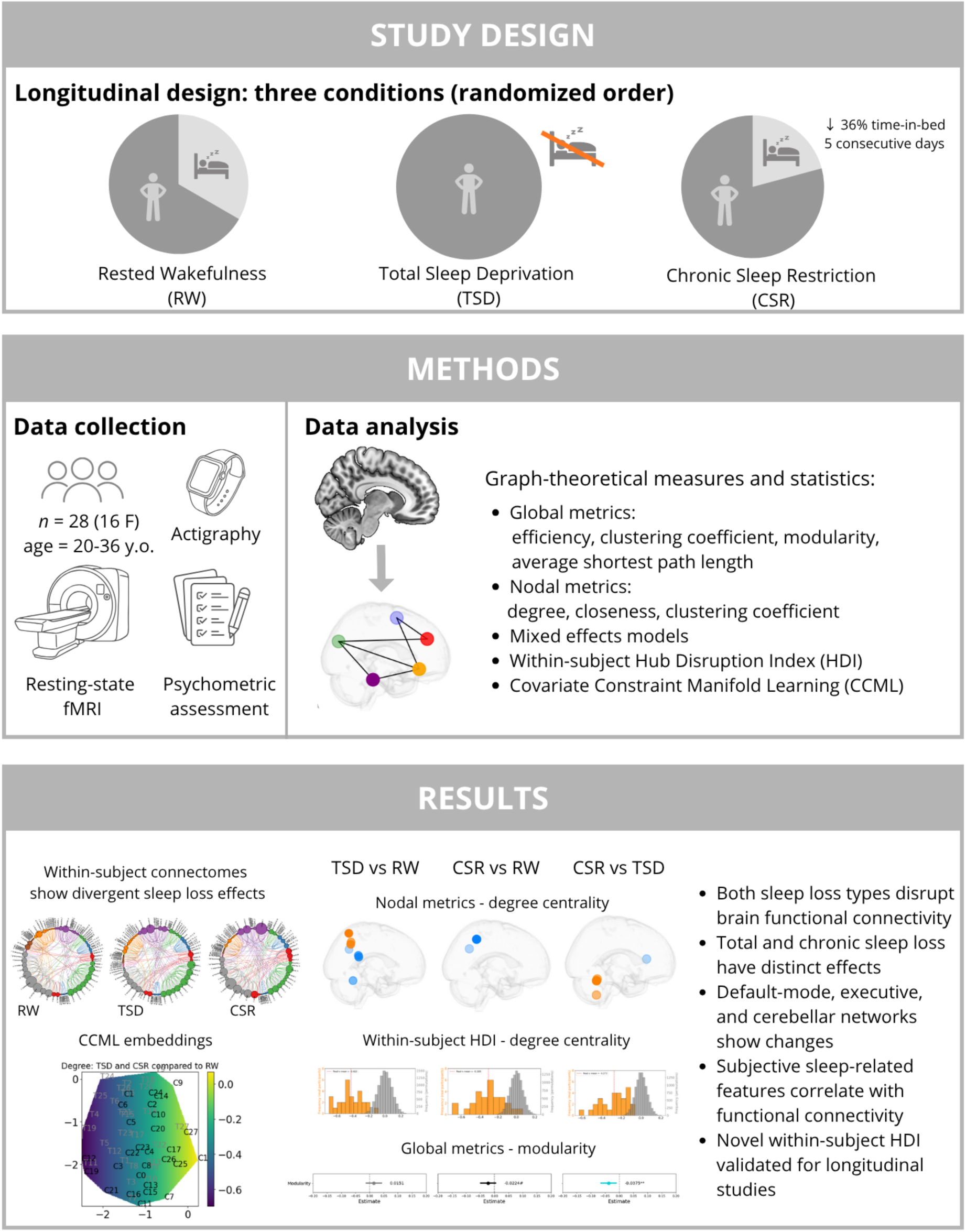

**Statement of significance:** Sleep deprivation is a growing public health concern, yet it remains unclear whether total and chronic sleep loss affect the brain in the same way. In this study, participants underwent one sleepless night and five consecutive days of reduced sleep (as in working week). We found that these two forms of sleep loss disrupt brain networks in fundamentally different ways. This distinction closes a critical gap in understanding how the brain responds to the total and chronic sleep loss. Recognizing that not all sleep deprivation is alike has important implications for managing fatigue in occupational, educational, and clinical contexts. Our findings highlight the need for personalized strategies to protect brain health and performance under conditions of sleep disruption.

## Introduction

Sleep is crucial for maintaining numerous aspects of human health and well-being. It is essential for cognitive abilities, including learning, memory consolidation, and executive functions [1–3]. Adequate sleep is also necessary for emotional stability and mental health [4,5]. Additionally, it plays a critical role in the regulation of various biochemical, metabolic, and immune processes [3,5].

Despite its fundamental importance, sleep is increasingly disrupted in modern society, with sleep disorders affecting a growing proportion of the population. In 2014, the U.S. Centers for Disease Control and Prevention described sleep deprivation as a “public health epidemic” linked to a wide range of medical issues, including hypertension, diabetes, depression, obesity and cancer [6–10].

Sleep deprivation is defined as the condition whereby an individual experiences insufficient quantity and/or quality (fragmentation) of sleep to support optimal alertness, performance, and health. This condition can be classified based on the degree and duration of sleep curtailment relative to normative sleep requirements. Acute Total Sleep Deprivation (TSD) refers to the elimination of sleep for a prolonged period of time (at least one night, without naps or other sleep episodes) [11]. Chronic sleep restriction (CSR) is defined as a consistent reduction in sleep duration below an individual’s habitual sleep need [12]. Both forms of sleep loss cause pronounced sleepiness, cognitive slowing, and mood changes in healthy adults [12,13].

A number of neuroimaging studies have shown that sleep loss disrupts functional networks [14–16]. For instance, acute TSD significantly decreases brain activity in the fronto-parietal attention network and salience network during cognitive tasks [15], and diminishes resting-state functional connectivity between the amygdala and the executive control areas [17]. Furthermore, the default mode network (DMN), which is involved in self-reflection, mind-wandering and mental state [18], is disrupted due to sleep deprivation [19]. The proper functioning of the brain requires the dynamic balance between “internally oriented” DMN and other, “externally focused”, task-positive networks [20]. TSD disturbs this balance, leading to a general loss of network segregation, whereby brain regions that normally function independently become abnormally coupled to each other [21]. CSR also impacts brain connectivity. Farahani and colleagues showed that global network properties were significantly changed as a result of a week of CSR. The connections were altered mostly across the limbic system, DMN, and visual network [14]. Although chronic sleep loss is the most common form of sleep loss in modern society, it received less experimental attention than widely studied TSD. Van Dongen and colleagues findings suggest that CSR has accumulative properties [22]. Thus, prolonged, moderate sleep loss may gradually degrade brain network organization, potentially reaching a similar level of disruption as seen after acute deprivation. However, direct comparisons between these two forms of sleep deprivation have never been studied before.

Over the past decades, neuroscientists have leveraged graph-theoretical methods on functional magnetic resonance imaging (fMRI) data to characterize functional connectivity in the brain [23–28]. By treating brain regions as nodes and their functional connections as edges, we can quantify changes in network architecture. This mathematical framework captures features of integration and segregation of networks that are not apparent through seed-based methods or Independent Component Analysis (ICA) [29,30] . They describe how strongly regions are connected, but do not consider the structure of the brain network as a whole. In contrast, graph theory describes the topological properties of the entire network and provides quantitative metrics of its organization. Network measures such as degree (i.e., the number of connections per node), clustering coefficient (i.e., how tightly connected a node’s neighbors are), and shortest path length (i.e., how efficiently information travels between distant brain regions) have been widely used to characterize brain organization.

In this study, we used resting-state functional magnetic resonance imaging (RS fMRI) to perform a direct, within-subject comparison across two different sleep deprivation states. We gathered data from 28 participants. Each participant underwent scanning in three conditions: during Rested Wakefulness (RW), following acute Total Sleep Deprivation (TSD), and after five consecutive nights of partial Chronic Sleep Restriction (CSR). Building on prior research, we anticipated differences in brain connectivity between both sleep-deprived states (TSD and CSR) and the well-rested control condition (RW). However, the primary aim of this study was to directly compare the neural effects of TSD and CSR. Specifically, we sought to evaluate two competing hypotheses: (I) CSR exerts cumulative effects on brain networks, ultimately leading to disruptions comparable to those observed after TSD, and (II) the two forms of sleep loss impact brain function via distinct neurobiological mechanisms. To quantify these effects, we employed graph theoretical analyses, which enable a comprehensive characterization of both global and local properties of the brain network topology. Based on Hub Disruption Index (HDI), we developed a new framework to characterize longitudinal disruption of networks. This approach allowed us to assess how sleep deprivation alters the brain’s functional architecture, providing insight into the organization of neural networks under different sleep loss conditions.

## Materials and methods

### Participants

Twenty-eight healthy volunteers (16 women and 12 men; aged 20-36 years, mean ± SD: 23.45 ± 3.71 years) participated in the study. All participants declared habitual sleep durations of 6.5-9 hours per night. Inclusion criteria required good subjective health, no history of neurological or psychiatric disorders, no ongoing medication use, and abstinence from alcohol, nicotine, and other psychoactive substances during the study. Any visual impairments had to be corrected with contact lenses. Left-handedness and incompatibilities with MRI safety protocol were additional exclusion criteria.

Each participant underwent three MRI sessions, each following a different sleep condition: (a) rested wakefulness (RW), defined as unrestricted sleep for at least one week prior to the scan (baseline); (b) total sleep deprivation (TSD), with no sleep on the preceding night, and (c) chronic partial sleep deprivation (CSR), involving restricted sleep (shortening by one third relative to RW) over five consecutive nights, simulating a “typical work-week”. The order of these conditions was counterbalanced across participants, with a minimum 14-day period between each session to mitigate carryover effects. To verify compliance with the sleep manipulation protocols, participants wore actigraphy-based motion loggers continuously during the week preceding each scanning session. In practice, RW was associated with an average sleep duration of 7 hours and 56 minutes (±44 min), CSR with 5 hours and 2 minutes (±17 min), representing a 36% reduction compared to RW, and TSD with no sleep (0 hours), corresponding to a 100% reduction. In TSD session, the total wakefulness duration prior to scanning was 26 hours and 5 minutes (±2 h and 23 min).

The procedure has been approved by the research ethics committee of the Institute of Applied Psychology at the Jagiellonian University (opinion No 126/2022). The study was conducted in accordance with ethical standards described in the Declaration of Helsinki. Every participant was informed about the procedures and goals of the study they volunteered in and gave their written consent. Participants were remunerated for taking part in the study.

### Scans acquisition

MRI data were acquired using a 3T scanner (Magnetom Skyra, Siemens) with a 64-channel head coil. High-resolution, whole-brain anatomical images were acquired using a T1-MPRAGE sequence (208 sagittal slices; voxel size 0.9 × 0.9 × 0.85 mm^3^; TR = 1800 ms, TE = 2.37 ms, flip angle = 8°). Functional T2*-weighted images were acquired using a whole-brain echo-planar (EPI) pulse sequence (48 axial slices, 3 × 3 × 3 mm isotropic voxels; TR = 1250 ms; TE = 27 ms; flip angle = 70°; MB acceleration factor 2; in-plane GRAPPA acceleration factor 2 and phase encoding A > P) using 2D multiband echo-planar imaging (EPI) sequences from the Center for Magnetic Resonance Research (CMRR), University of Minnesota [31–33]. Resting state scan lasted 8 minutes.

The study was conducted in the Malopolska Centre of Biotechnology at the Jagiellonian University in Kraków. All the registrations were carried out between 9:30 a.m. and 3:30 p.m. Regardless of the weather conditions outside, the temperature, lighting and noise levels in the laboratory were constant.

### Preprocessing

Preprocessing of fMRI images was performed using custom MATLAB (R2022b) scripts built on top of SPM12 (Wellcome Trust Centre for Neuroimaging, University College London, UK), Computational Anatomy Toolbox, CAT12 [34], and the CONN toolbox [35,36].

For each participant, preprocessing of the structural T1-weighted MRI image consisted of the following steps: (I) magnetic field inhomogeneity correction (bias regularization = 0.5), (II) segmentation into gray matter (GM), white matter (WM), and nonbrain tissues, resulting in tissue probability maps used for weighted time series extraction, (III) affine registration to MNI152 standard space, and (IV) nonlinear normalization using geodesic shooting as implemented in the CAT12 within SPM12.

Functional scans were corrected for head motion by realigning all volumes in each time series to the mean image. The mean resting state (RS) image was then coregistered with each subject’s T1-weighted structural MRI. This enabled the computation of the deformation field in two-step normalization: (I) mean functional image to T1-weighted structural space, and (II) T1-weighted structural image to MNI standard space. Based on that, the inverse of the estimated deformation field was calculated, allowing the MNI-space anatomical atlas and the gray matter (GM) probability map to be warped into each subject’s native RS functional space. The data were not spatially smoothed. This approach minimizes fMRI data distortions and ensures precise, individualized parcellation defined by the modified Anatomic-Automatic Labeling (AAL) atlas [37]. The classical AAL atlas was modified by merging some of the smallest regions into bigger structures, reducing the parcellation to 89 regions. Based on previous work, this modified atlas provides a suitable balance between anatomical precision, region size, and the resulting number of possible edges in the graph representations. Details of the modified atlas, are described in Supplementary materials S1 and in Achard’s, Termenon’s works [38,39].

Additionally, to compare the results obtained using the modified AAL atlas (an anatomically based parcellation), we reconducted all analyses using the AICHA atlas, which was constructed based on the functional connectivity patterns [40]. Since the AICHA atlas does not include the cerebellum, we extended it by incorporating time series from cerebellar parcellation based on the AAL atlas. This combined atlas comprises a total of 391 brain regions. Results obtained using this parcellation are presented in the Supplementary materials S12.

### Time series extraction

At each time point, in each parcel, regional mean time series were estimated by averaging the fMRI voxel values weighted by the GM probability maps (proportion of gray matter in each voxel of the segmented structural MRIs) [24], which reduces the noise in the time series caused by white matter and cerebrospinal fluids signals.

To diminish motion-related artifacts, nuisance regression was performed using regressors derived from the Artifact Detection Toolbox (ART) (https://www.nitrc.org/projects/artifact_detect/), including six motion parameters, their first-order temporal derivatives, and binary regressors representing outlier volumes. We did not perform despiking. Mean values were regressed out from the resulting time series to center them at zero.

We assessed the quality of the fMRI data, by calculating the Framewise displacement (FD) for each ROI in each time point and frame-by-frame percentage changes in BOLD signal throughout the brain across all participants (following the approach of Power and colleagues [41]). Quality check is shown in the Supplementary Figure S2.

The obtained set of regional mean time series was then used for wavelet correlation analysis and graph construction.

### Wavelet decomposition

Wavelet decomposition correlation analysis filters fMRI signals into time-frequency components using compactly supported basis functions, or little waves, each of which is uniquely scaled in frequency and located in time. This method is well suited for analyzing RS fMRI data, which often exhibit fractal or 1/f properties (for a review of wavelet methods for fMRI data analysis, see: [42–44]).

In order to perform the decomposition of the fMRI signal, we applied the maximal overlap discrete wavelet transform (MODWT) implemented in the *brainwaver* R package, to each regional mean time series (*N* = 89 for modified AAL atlas and *N* = 391 for modified AICHA atlas). Then, we estimated pairwise interregional correlations at each of the 4 wavelet frequency scales. We further used the wavelet scale 3 (which corresponds to the frequency range ∼0.05-0.1 Hz) since 0.1 Hz is commonly used as a threshold value to capture the relevant information in resting-state functional connectivity and these oscillations fall within the range where the small-world topology of the brain is the most salient [23,45]. The rationale for selecting this wavelet scale and frequency bands is detailed in the corresponding articles [24,46].

### Graph construction

For each participant, functional connectivity was quantified by computing pairwise Pearson correlation coefficients between all 89 regions of interest (ROIs), resulting in an 89×89 correlation matrix. In this study, we binarized correlation matrices derived from the modified AAL atlas by retaining 200, 400 and 600 strongest correlations. These values correspond to approximately 5%, 10% and 15% graph cost, respectively, where graph cost is defined as the ratio of the number of retained edges to the total number of possible edges in the graph (here: 3916 possible edges) [38,47]. To ensure comparability across participants, we kept the number of edges in each graph fixed. To keep graphs fully connected, a minimum spanning tree [48,49] based on the distance transformation of the absolute correlation values was applied prior to thresholding. This process results in an adjacency matrix, which defines the unweighted, undirected, fully connected graph with a fixed number of binarized edges for each subject.

In this article, we present results based on graphs constructed using 400 edges (corresponding to a 10% graph cost), as this threshold, in our experience, yields networks with a balanced level of sparsity and density. To assess the robustness of our findings, additional results using 200 and 600 edges (representing 5% and 15% graph costs, respectively) are provided in Supplementary Materials S11.

These additional calculations were motivated by the fact that a gold standard for determining the optimal number of edges and thresholding strategy is still lacking. The lower limit of edges is defined by the minimal number of edges required for the graph to diverge meaningfully from a Erdos-Renyi or regular graphs. The upper limit is reached when edge density becomes so high that the graph’s structure is no longer informative, approaching the properties of a random or complete graph. The range of edge densities is also constrained by statistical significance: a minimum number of significant edges is required to ensure structural coherence, while excessive inclusion of weak or non-significant correlations may inflate edge density and obscure meaningful topological patterns [46,50]. We provide the number of significant edges for each participant in each scanning session in Supplementary materials S3.

For the modified AICHA atlas that consists of 391 nodes, we followed the same graph construction procedure using 2200 edges, which corresponds to a 3% graph cost. This threshold was selected as the lowest common number of possible edges across all 28 participants along their three scanning sessions. This approach allowed us to retain the full cohort without excluding any participants due to insufficient edge count. The number of significant edges and the corresponding results using these graphs are also presented in Supplementary Materials S3, S12.

Wavelet-based functional connectivity analysis and graph construction were carried out using the brainwaver and igraph R libraries. Detailed description of this procedure is available in the Termenon and colleagues work [47].

### Graphs metrics

To measure the topological features of graphs, we used both nodal and global metrics.

For each node of each individual graph, we calculated three commonly used graph metrics: degree centrality, clustering coefficient and closeness centrality. Degree is defined by number of edges connected to a node *i*. Degree centrality is normalized by dividing degree value by the highest node degree in the network. We also employed the clustering coefficient, which indicates how interconnected the neighbors of node *i* are, providing insight into local network density. Finally, closeness centrality, which captures how efficiently information can spread from a node *i* to the rest of the network. Together, these metrics offer a three-level assessment of graph structure: the node itself (degree centrality), its local neighborhood (clustering coefficient), and its global positioning in the network (closeness centrality). The formulas for calculating these three metrics are provided in the supplementary material S4.

Moreover, to further characterize the connectivity graphs, we computed the global metrics, which are calculated across all nodes in each network, resulting in the single value per graph, instead of single value per node. Schematic comparison between nodal and global metrics (using nodal clustering coefficient and global average clustering coefficient as example) is presented in Figure 1 We focused on the global efficiency, average clustering, average shortest path length, modularity, and average graph distance between communities. Detailed descriptions of all these metrics can be found in [28,51,52].

**Figure 1.**
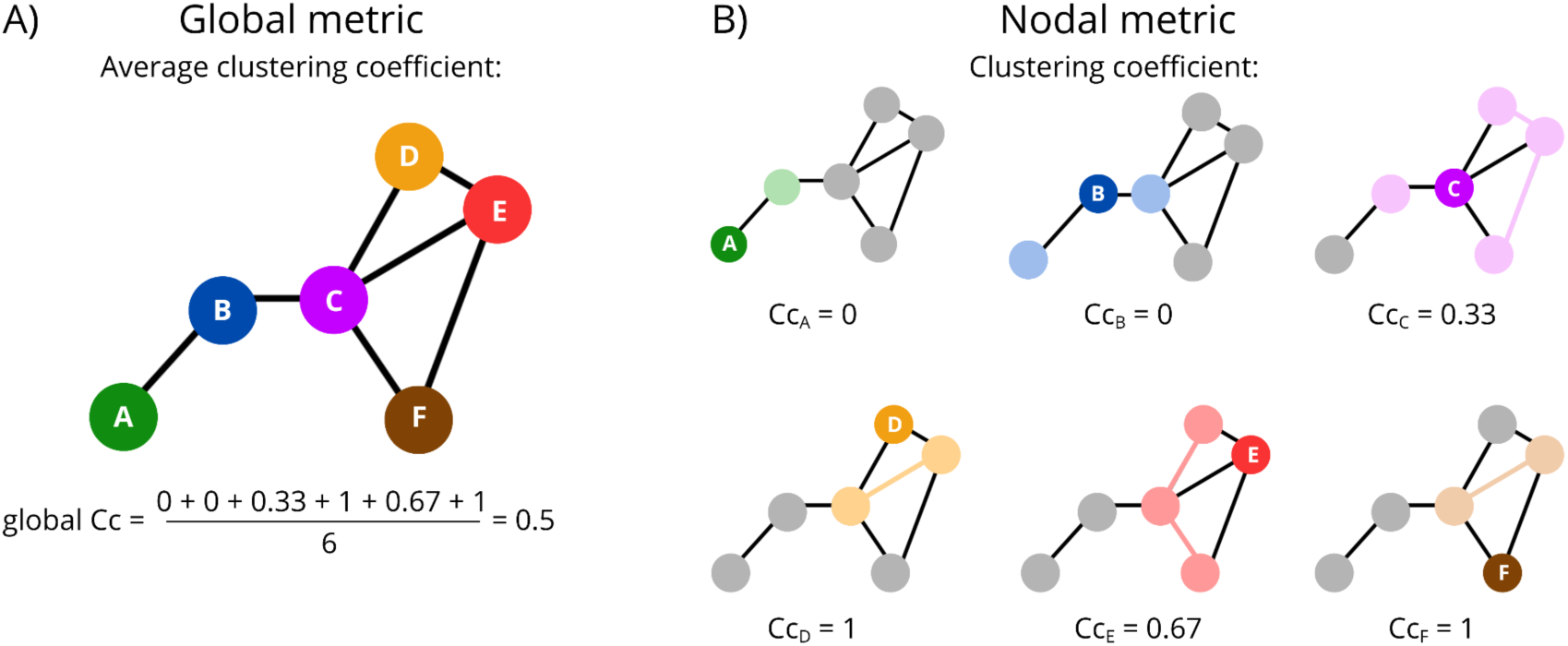
Schematic representation of global and nodal metrics, using the clustering coefficient (Cc) as an example. A consistent color code is used to preserve node identity across all panels. A) Global clustering coefficient, calculated as the mean of all nodal clustering coefficient values. B) Nodal clustering coefficients. For each node, its direct neighbors and the connections between them are marked using lighter shades of the corresponding node color.

### Statistical analyses

#### Global and nodal metrics comparisons

Global and nodal graph metrics were compared between experimental conditions using paired t-tests and linear mixed-effects models (LMM) to account for both within-subject and between-subject variability. This approach was chosen due to the non-independence of observations, as each participant was scanned three times under different sleep conditions. Correction for multiple comparisons was performed using the False Discovery Rate (FDR) approach and the Benjamini-Hochberg linear step-up (LSU) procedure, with a false discovery rate threshold of q<0.05 for global metrics and q<0.1 for nodal metrics. To assess the robustness of findings, we conducted two types of non-parametric permutation tests with 10,000 iterations. First, a paired permutations approach was used, in which we preserved within-subject pairs but randomly mixed assigned session labels. Second, unpaired permutation test was used, in which we shuffled all values across sessions without preserving subject identity, thus breaking the within-subject pairing.

All analyses were performed using Python 3.9.23 and NetworkX library 3.2.1 [53] for graph analysis, NumPy 1.26.4, Pandas 2.2.3 for data manipulation, Matplotlib 3.9.2, Seaborn 0.13.2, Netplotbrain 0.3.0, and HoloViews with Bokeh backend for visualization, and SciPy 1.12.0, Statsmodels 0.14.4 for statistical testing.

#### Hub Disruption Index

The Hub Disruption Index (HDI) is a metric designed to evaluate the disruption of network topology of a given subject (e.g. patient) relative to a reference network structure [24,47]. It quantifies this reorganization by comparing node-level metrics (e.g. degree centrality) across all brain regions to the corresponding averaged values obtained from healthy control group. Thus, it is a global index capturing changes at the nodal level.

Given our longitudinal dataset, which enables within-subject comparisons, we adapted this method by using each participant’s own baseline scan as a reference, rather than the mean of the healthy control group baseline. It is well-known that healthy volunteers show high interindividual variability [54]. In this work, we introduce a new way to avoid these spurious effects.

The within-subject HDI calculation involves three key steps: (I) computation of the node-level metric (e.g. degree centrality) for each subject in all conditions, (II) computation, for each node, the difference between the nodal metric in condition-of-interest and nodal metric in reference condition, (III) plotting the difference values for each node against the nodal metric from reference condition, then fitting a linear regression. The slope of regression line represents the 𝜅 value from HDI. This procedure has to be repeated for each participant and for each conditions comparison, resulting in a set of 𝜅 values.

For a formal definition, let *N* represent the number of nodes in the graph, *n* the number of participants, and *T* the number of experimental conditions. For each subject *s*, we define a nodal metric matrix 𝑀^(*s*)^ ∈ 𝑅^*N✕T*^, where each element *m_i_^(s,t)^* represents the graph metric of node *i* in condition *t*. Then, the within-subject HDI is defined by the linear model:

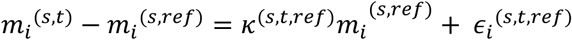

where *mi^(s,t)^* represents nodal graph metrics of node *i*, for subject *s* in the condition-of-interest *t*, *mi^(s,ref)^* represents nodal graph metrics of node *i*, for subject *s* in the reference condition, 𝜅*^(s,t,ref)^* is the HDI slope for subject *s*, comparison *t vs ref*, describing the graph disruption, and 𝜖_*i*_*^(s,t,ref)^*is the classical error term in linear regression.

A 𝜅 value close to zero indicates no systematic difference in network topology between the examined conditions: graph metric obtained at the node level is the same for the condition-of-interest and the reference condition. In contrast, positive or negative 𝜅 values indicate a reorganization of hubness, with some regions becoming more central (increased hubness) and others less central (decreased hubness), reflecting reorganized network topology.

In our study, each participant was scanned in three conditions, thus the relative difference in the functional connectivity of the same brain between any two states was calculated (Figure 2). This approach enables us to capture the relative, subject-specific disruption in graph architecture between the sleep-deprived and a control state, effectively tailoring the HDI to the repeated-measures nature of our study.

**Figure 2.**
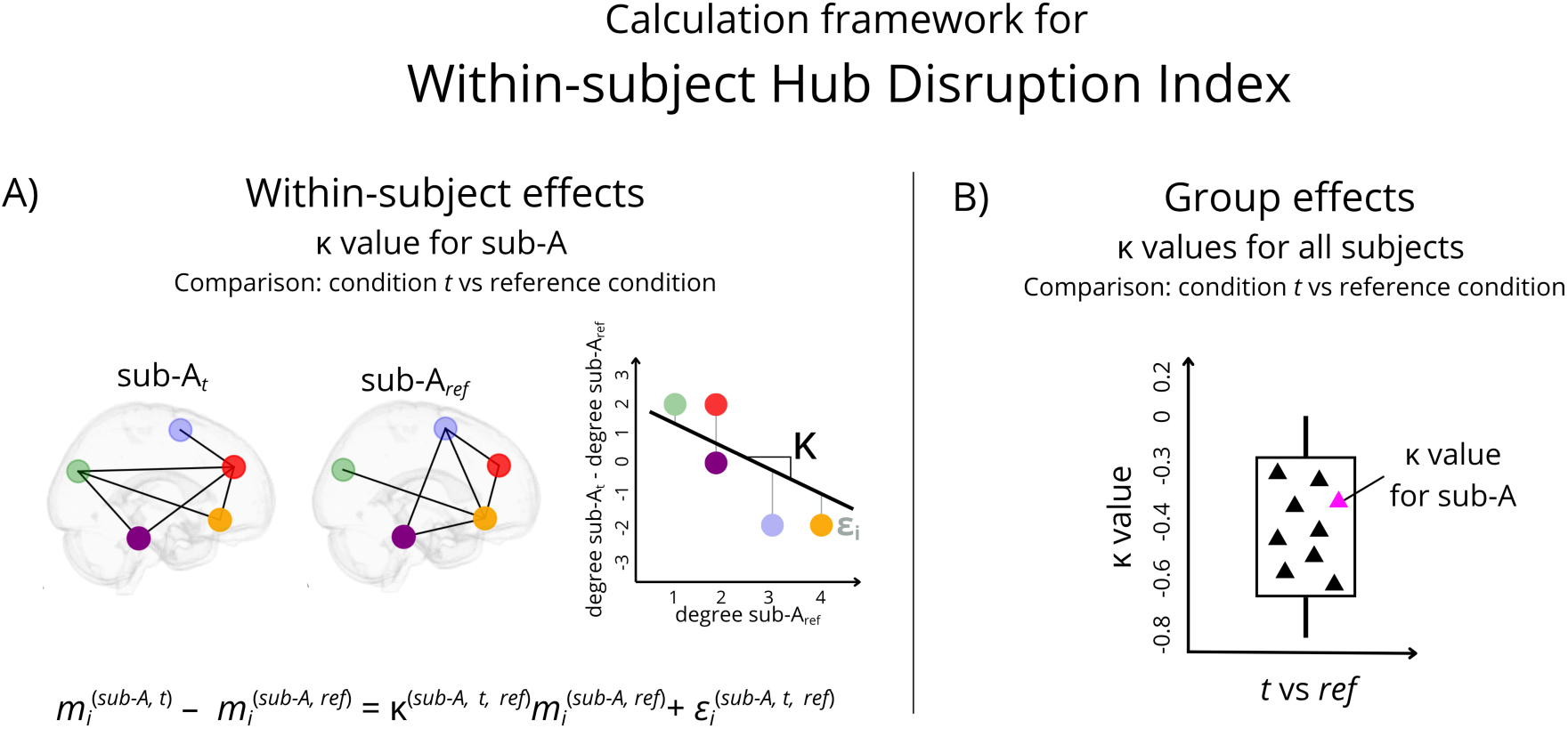
Calculation framework for within-subject Hub Disruption Index (HDI). This figure illustrates the within-subject computation of the HDI using nodal metric (*m_i_*, here: degree) for a representative participant (sub-A) for comparison: condition *t* vs reference condition. In this study, we conducted three comparisons: TSD vs RW, CSR vs RW, and CSR vs TSD. A) Schematic representations of the individual κ value calculation. The scheme contains simplified brain graphs with five nodes and six edges for both, condition *t* and reference condition. A scatter plot displays degree values from the reference condition (x-axis) against the difference in degree between condition *t* and the reference (y-axis). Node identities are preserved across conditions (via consistent colors) to illustrate network disruption. A linear regression model is fitted to the data, yielding a κ value (slope of regression line). B) Schematic representation of group-level effects, illustrated as a boxplot of κ values across all participants for a given comparison (conditions *t* vs ref). The κ value of sub-A is highlighted in magenta. Abbreviations: RW - rested wakefulness, TSD - total sleep deprivation, CSR - chronic sleep restriction,

#### Permutations

The within-subject HDI is designed for the repeated-measures of our longitudinal dataset. We further validated the robustness of this approach by using three different, complementary permutation tests, as described below. For each test, we used 10,000 permutations to generate a null distribution to compare it against our observed values and test for statistical significance. Schematic illustration of our three approaches is shown in Supplementary material S5.

##### First permutation test

We aimed to evaluate whether the within-subject HDI (𝜅_within_) is more sensitive to individual disruption patterns than the group-level HDI (𝜅_group_) and check if the difference between within-subject and group-level HDI is a result of the random anatomical correspondence.

We constructed a null distribution by shuffling the node labels in the graph corresponding to the sleep deprivation condition, while preserving the node labels in the control condition and maintaining the original subject-level pairing and conditions labels.

Then we calculated HDI using two approaches - within subject and group-level. We computed 𝜅_within_ for each participant by comparing their individual reference graph to their permuted sleep-deprived graph. In parallel, we calculated 𝜅_group_ by comparing each participant’s permuted sleep-deprived graph to the group-averaged degree values from the reference condition. The same regression procedure was applied, yielding both, 𝜅_within_ and 𝜅_group_ values per participant.

Finally, for each permutation, we calculated the difference between 𝜅_within_ and 𝜅_group_. A negative difference (i.e. 𝜅_within_ - 𝜅_group_) would indicate that stronger hub disruption is detected when using the subject’s own baseline as a reference, compared to the group average. Because the node labels were shuffled in the sleep-deprived graph, this test specifically evaluates whether the added sensitivity of the within-subject HDI depends on preserving spatial (anatomical) correspondence between sessions. The negative value would support the hypothesis that the within-subject HDI captures subject- and node-specific disruption patterns that may be diluted in the group-level approach.

##### Second permutation test

Given the longitudinal design of our dataset, we tested whether subject-level pairing contributes to the sensitivity of the within-subject HDI. To examine this, we randomly mixed the order of scans, breaking the original subject pairing and comparing each control with a sleep deprived scan from a different subject. A null distribution centered in values more negative than those observed would imply that comparing graphs coming from different brains, rather than the same brain across conditions, introduces greater variability leading to spurious disruption measures.

##### Third permutation test

We aimed to test whether the observed differences between conditions could be explained by random condition assignments. To this end, we designed a permutation test in which the condition labels were randomly switched within each subject, while preserving subject-level pairing. For each permutation, we computed the within-subject HDI and compared the results to those obtained from the original data with the correct labels.

Results of these procedures are presented in Supplementary Figure S6. These three permutation tests allow us to challenge our findings against randomness. These are empirical tests, however, we think that these tests highlight the robustness of our proposed approach.

### Covariate-Constraint Manifold Learning

In addition to the classical nodal- and global-level metrics, we applied Covariate-Constrained Manifold Learning (CCML) - a dimensionality reduction technique, designed specifically for neuroimaging and network neuroscience applications, recently introduced by Renard and colleagues [55]. The goal of applying CCML was to identify low-dimensional representations of individual functional network topologies and classify them to the different sleep deprivation states.

It has already been shown that global metrics may not discriminate between two groups of subjects [24], which highlight their limits as biomarkers. While nodal-level approaches offer better spatial specificity, they also pose their own challenges: hundreds of brain regions can be extracted, whereas the number of subjects is typically small. This leads to the high-dimension, low-sample-size (HDLSS) scenario - configuration in which the number of parameters far exceeds the number of observations. Such data are particularly vulnerable to the *curse of dimensionality* [55,56], which can compromise the robustness of standard classification and regression algorithms [57]. Thus, manifold learning provides a framework to reduce complexity while preserving biologically meaningful variation in network data. However, traditional unsupervised manifold learning methods (e.g., ISOMAP) are purely data-driven and do not incorporate any external covariates, which can limit the interpretability of the resulting embeddings. As an extension of this approach, CCML constrains one coordinate of the reduced space to correspond to a specific, biologically meaningful covariate. The other coordinates are left unconstrained, as in classical ISOMAP.

To apply CCML in our study, each subject was represented by a combination of their κ value (used as the covariate) and a nodal feature vector derived from degree centrality, forming the input for dimensionality reduction.

The reduced point *x̃*_i_ is defined by *x̃*_i_ = [*αc*_i_; *x*_i_]*^T^*, where *c*_i_ is the chosen covariate (in this study: κ value) and *x*_i_ represents the other coordinates in the embedding space. A scaling factor *α* is introduced to balance the covariate axis with the rest of the dimensions. Factor *α* is estimated by optimization. We initialized the *α* parameter based on the relative spread of κ values and the initial ISOMAP embedding. To improve embedding stability, we selected the first two subjects as the most distant pair based on ISOMAP geodesic distances and then iteratively added the subject that was farthest from those already embedded. CCML then sequentially optimized each subject’s position in embedding by minimizing the discrepancy between geodesic distance and the distance in combined network-covariate space, followed by global refinement of both subject coordinates and the *α* scaling parameter. Detailed description of CCML optimization strategy can be found in [55]

The final embedding was visualized in 2D, combining κ values and nodal metric (degree centrality), allowing us to assess group separability and network disruptions associated with TSD and CSR.

Manifold learning and dimensionality reduction were performed using scikit-learn, and nonlinear optimization was carried out with SciPy’s optimize module in Python.

### Behavioral measures

In this study, we distinguished between two categories of subjective sleep-related features: state-level features, reflecting the participant’s current subjective condition, and trait-level features, reflecting stable individual characteristics. To quantify them, we employed the following psychometric questionnaires: Karolinska Sleepiness Scale (KSS [58]), Pittsburgh Sleep Quality Index (PSQI [59]), Oginska Chronotype Questionnaire [60], with two subscales for morningness-eveningness, ME, and subjective amplitude of the circadian rhythm, AM. The ME represents an individual’s preferred time of activity, while the AM - the strength of this preference, understood as the subjective amplitude between hyper- and hypo-activation phases [60–62]. Extreme chronotypes were excluded from our cohort.

KSS data was collected during each experimental session (representing subjective sleepiness in RW, TSD and CSR), while PSQI, ME, AM were collected once, before start of the experiment.

### Behavioral data analysis

We defined 3 research questions regarding subjective behavioral measurements: (I) How is subjective sleepiness (state-level) related to objective graph-theoretical measures in each session (RW, TSD, CSR)? (II) Which subjective trait-level measures predict brain functional networks metrics in baseline RW condition? (III) Which subjective trait-level measures predict the reorganization of functional networks following sleep deprivation?

To address first question, we employed the linear Mixed Effect Model. Specifically, we tested whether subjective sleepiness was associated with global graph metrics (global efficiency, average clustering coefficient, average path length, modularity, and average graph distance) within each scanning session (fixed effect) while accounting for repeated measurements per participant (random effect). We controlled the false discovery rate (FDR) within each metric using the Benjamini-Hochberg procedure.

To examine whether trait-level sleep and circadian measures predict brain global graph metrics, first, we confirmed no collinearity between PSQI, ME and AM and z-scored these values to enable comparability of effect sizes across measures with different scales. Then, we fitted the ordinary least squares (OLS) regression models with HC3 heteroskedasticity-consistent standard errors. For each global graph metric in the RW session, all three trait measures were simultaneously included as predictors.

To test whether traits (AM, ME, PSQI) predict network disruptions (defined as κ values from HDI) we used analogous OLS regression models. For each κ outcome for three nodal metrics (degree centrality, closeness centrality, clustering coefficient), all three traits were entered simultaneously as predictors. For both types models, p-values were adjusted using the Benjamini-Hochberg FDR procedure (*q* = 0.05).

All three predictors were not correlated with each other.

## Results

### Global graphs metrics

Using linear mixed-effects models in which sleep deprivation conditions were included as fixed effects, and within-subject variability was modeled using subject ID as a random effect, we found that CSR is related to decreased value of modularity in comparison to TSD (*β* = -0.0375, *p* = 0.0276). Differences in all other metrics are not significant as *p* > 0.05. Results are shown in Figure 3, detailed information can be found in the Supplementary Table S7.

**Figure 3.**
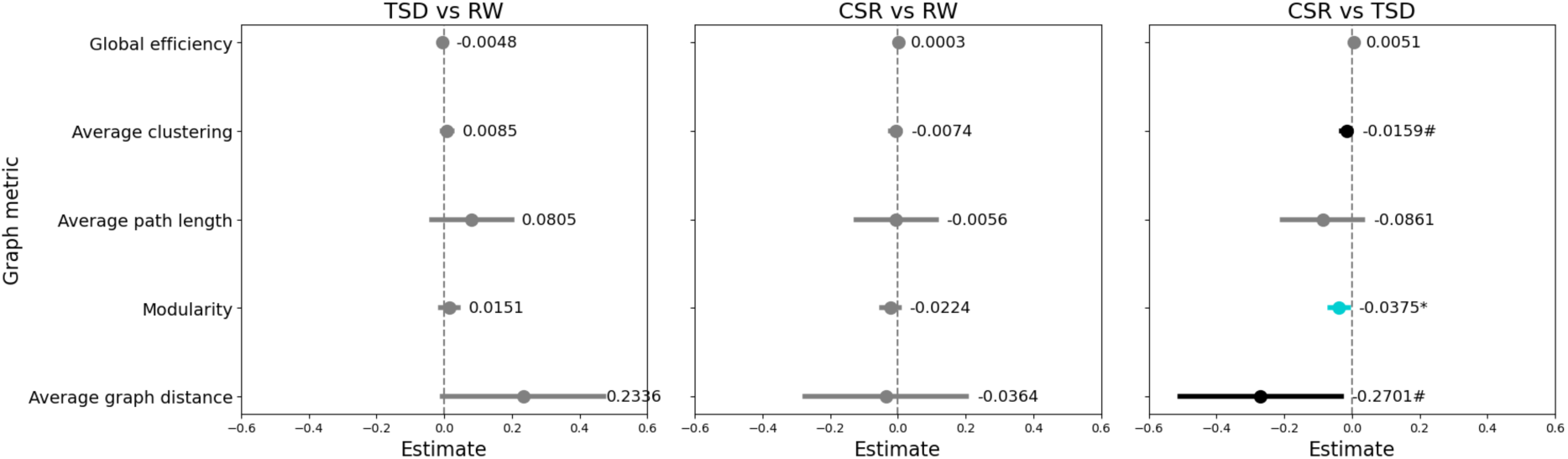
Linear mixed-effects model (LMM) regression results for global efficiency, average clustering, average path length, modularity, and average graph distance between communities. The left panel shows the comparison TSD vs RW, the middle panel shows CSR vs RW, and the right panel shows CSR vs TSD (with TSD treated as the reference). Dots and numerical labels represent *β* coefficient estimates, horizontal bars show confidence intervals (95% CI). Statistically significant association (after FDR multiple comparisons correction) is marked in blue, * represents *p*-value ≤ 0.05, trend-level associations are marked in black, # represents *p*-value ≤ 0.1. Abbreviations: RW - Rested wakefulness, TSD - Total Sleep Deprivation, CSR - Chronic Sleep Restriction.

To check the robustness of these results, we conducted two types of non-parametric permutation-based (10,000 iterations) paired t-tests for each session pair (TSD vs RW, CSR vs RW, CSR vs TSD). These robustness checks are reported in the Supplement Figure S8.

Reported results indicate that functional connectivity, as reflected by changes in global graph metrics, is altered in distinct ways between the two sleep deprivation conditions. No significant changes were observed when either sleep deprivation condition was compared to the control state. This suggests that averaging used to obtain global metrics may overly smooth the results, highlighting the need for further investigation at the graph nodal level.

### Nodal graph metrics

Degree centrality, closeness centrality and clustering coefficient calculated for each subject in each condition and compared between conditions using linear-mixed effects model revealed changes between both sleep deprivation states and RW, as well as between two sleep deprivation types. Figure 4 and Table 1 show details about observed changes.

**Figure 4.**
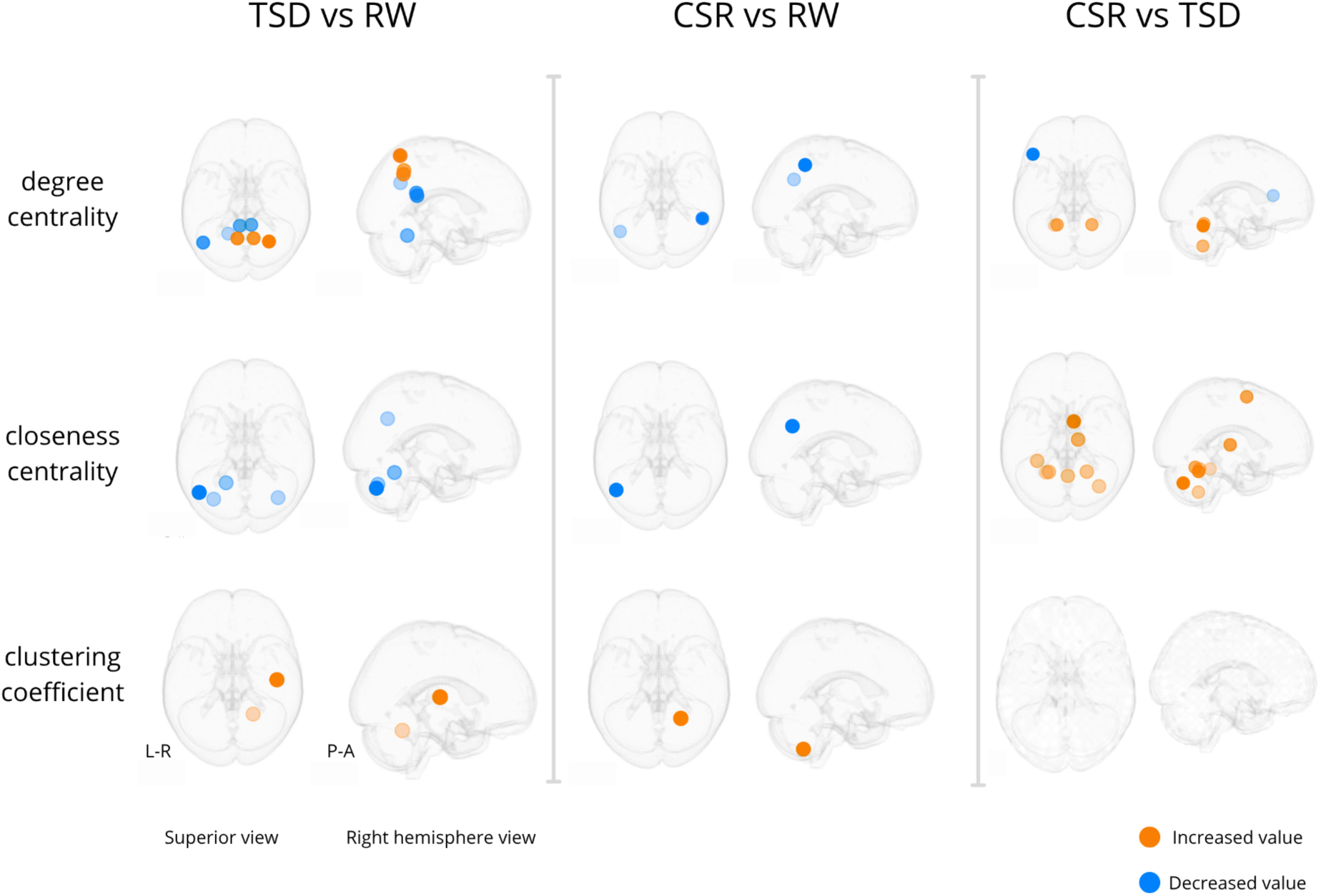
Brain regions showing significant changes in nodal metrics (degree centrality, closeness centrality, and clustering coefficient) across different sleep deprivation states. Orange dots indicate regions with increased metric values, blue dots indicate decreased values. Node locations are defined based on centroids of regions from the modified AAL atlas (89 brain regions). Statistical significance was determined using the linear mixed-effects model followed by multiple comparisons correction (FDR correction, *q* < 0.1). Visualization was prepared using the Netplotbrain Python package [63] and overlaid on the MNI glass brain template. Abbreviations: RW - Rested wakefulness, TSD - Total Sleep Deprivation, CSR - Chronic Sleep Restriction.

**Table 1.** Brain regions showing significant changes in nodal metrics across sleep deprivation states. Each row presents the anatomical region (based on the modified AAL atlas), the centroid MNI coordinates, and the direction and magnitude of changes in nodal metrics (degree centrality, closeness centrality, clustering coefficient) for each pairwise session comparison: TSD vs RW, CSR vs RW, and CSR vs TSD. Arrows indicate the direction of change (↑ = increased value, ↓ = decreased value), and values in parentheses represent the corresponding coefficients from the linear mixed-effects model. Only regions with significant changes after FDR correction (*q* < 0.1) are included. Abbreviations: RW - Rested wakefulness, TSD - Total Sleep Deprivation, CSR - Chronic Sleep Restriction.

| Region | MNI Coordinates | TSD vs RW | CSR vs RW | CSR vs TSD |
| --- | --- | --- | --- | --- |
| Left Angular Gyrus | (-45.3, -62.3, 33.0) | Degree ↓ (-0.0260)<br>Closeness ↓ (-0.0311) | Degree ↓ (-0.0369)<br>Closeness ↓ (-0.0346) | |
| Left Cerebellum Lobules III–VI | (-19.3, -54.3, -22.6) | Degree ↓ (-0.0702)<br>Closeness ↓ (-0.0508) | | Degree ↑ (0.0929)<br>Closeness ↑ (0.0566) |
| Right Cerebellum Lobules III–VI | (21.7, -54.1, -24.0) | Clustering ↑ (0.1338) | | Degree ↑ (0.0828)<br>Closeness ↑ (0.0586) |
| Left Cerebellum Lobules I–II | (-32.6, -71.9, -35.2) | Closeness ↓ (-0.0422) | | |
| Right Cerebellum Lobules I–II | (35.3, -70.4, -36.2) | Closeness ↓ (-0.0554) | | Closeness ↑ (0.0438) |
| Left Cerebellum Lobules VII–X | (-22.7, -55.7, -49.2) | | | Degree ↑ (0.0649)<br>Closeness ↑ (0.0523) |
| Right Cerebellum Lobules VII–X | (23.3, -57.5, -50.6) | | Clustering ↑ (0.1403) | |
| Left Posterior Cingulate Cortex | (-6.0, -44.5, 22.0) | Degree ↓ (-0.0239) | | |
| Right Posterior Cingulate Cortex | (6.4, -43.5, 19.2) | Degree ↓ (-0.0272) | | |
| Left Inferior Frontal Gyrus (Triangular Part) | (-46.7, 28.1, 10.1) | | | Degree ↓ (-0.0353) |
| Left Fusiform Gyrus | (-31.9, -42.4, -23.1) | | | Closeness ↑ (0.0274) |
| Right Heschl's Gyrus | (45.1, -18.6, 7.7) | Clustering ↑ (0.3292) | | |
| Right Inferior Parietal Lobule | (45.3, -47.5, 47.2) | | Degree ↓ (-0.0244) | |
| Right Superior Parietal Lobule | (24.8, -60.2, 59.8) | Degree ↑ (0.0402) | | |
| Left Precuneus | (-8.5, -57.4, 45.5) | Degree ↑ (0.0463) | | |
| Right Precuneus | (8.8, -57.4, 41.3) | Degree ↑ (0.0463) | | |
| Right Supplementary Motor Area | (7.2, -0.9, 58.7) | | | Closeness ↑ (0.0347) |
| Right Thalamus | (12.0, -19.3, 5.2) | | | Closeness ↑ (0.0428) |
| Cerebellar Vermis | (1.2, -58.6, -20.1) | | | Closeness ↑ (0.0475) |

The comparison between TSD and RW showed that TSD primarily affects the posterior and parietal regions of the Default Mode Network (DMN), which is observed as change in degree centrality value. We observed a dissociation between DMN subsystems. Regions constituting the Dorsal Medial subsystem (such as the Left Angular Gyrus and the bilateral Posterior Cingulate Cortex) show a decrease in connectivity. Additionally, reduced degree is seen in the sensorimotor region of the left Cerebellum (lobules III–VI). In contrast, regions associated with another DMN subsystem - specifically the Right Superior Parietal Lobule and the bilateral Precuneus - exhibit increased degree. Moreover, an increase in clustering coefficient in the Right Heschl’s Gyrus and Right Cerebellum Lobules III–VI suggests that the neighboring nodes of these regions have become more strongly interconnected.

We found that CSR also leads to a reduction in connectivity within DMN regions, including the Left Angular Gyrus and the Right Inferior Parietal Lobule.

In contrast, the comparison between two types of sleep deprivation demonstrated that CSR, relative to TSD, results in decreased connectivity in the Anterior part of the Frontoparietal network (FPN), specifically in the Left Inferior Frontal Gyrus (triangular part). At the same time, increased connectivity is observed in the cerebellum: in both left and right cerebellar lobules III-VI (sensorimotor regions) and in the left cerebellar lobules VII-X (cognitive and limbic regions). Additionally, increase in closeness centrality in the Right Supplementary Motor Area, Right Thalamus, and Cerebellar Vermis suggests that these regions become more globally accessible and functionally integrated, potentially serving as hubs that facilitate efficient information transfer.

#### Hub Disruption Index

To further explore the changes in the topology of functional connectivity graphs, we employed the Hub Disruption Index (HDI), adapted for longitudinal data (within-subject HDI).

Consistently negative κ values were observed across all subjects and for all three graph metrics (degree centrality, closeness centrality, clustering coefficient) when comparing both sleep-deprived conditions to the RW. Furthermore, comparisons between TSD and CSR conditions also yielded uniformly negative κ values, suggesting that these two sleep deprivation types differentially affect network topology, despite both showing a general trend toward disruption.

To confirm the robustness of our results, we used permutation tests in which the condition labels were randomly shuffled within each subject, while preserving subject-level pairing. For each permutation, we computed the within-subject HDI and compared the results to those obtained from the original data with the correct labels. Cohen’s *d* values ranged from -3.8 to - 5.1) and two-tailed *p*-values from 10,000 permutations were < 0.001 after FDR correction, which confirms the robustness and statistical significance of the observed effects. Full distributions of κ values and permutation-based null distributions are presented in Figure 5.

**Figure 5.**
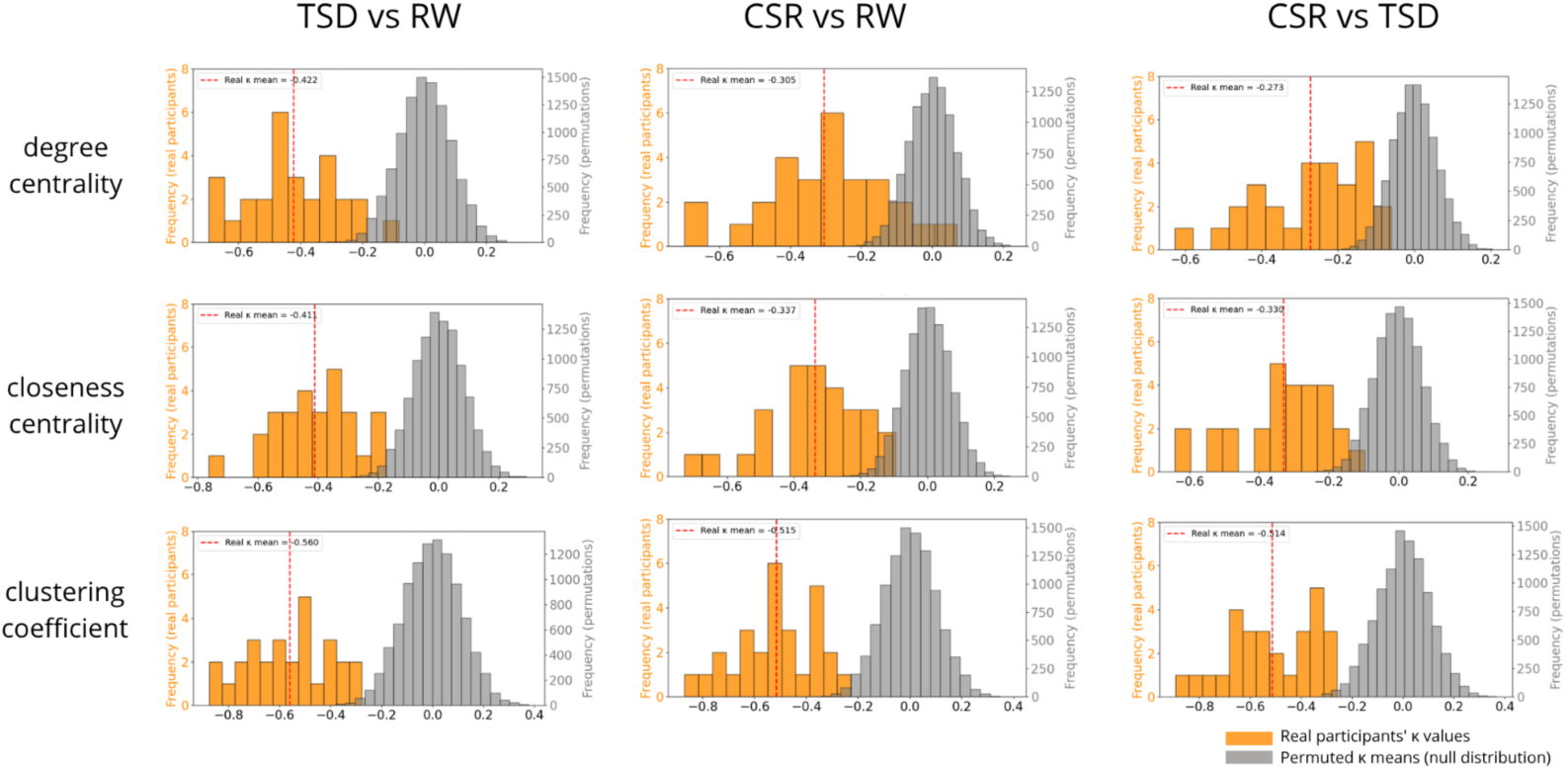
Result of within-subject Hub Disruption Index (HDI) analysis. Orange distributions represent κ values obtained from each participant for each comparison and nodal metric (degree centrality, closeness centrality, clustering coefficient). The red dashed line indicates the group mean κ value for each comparison. The grey distribution shows the null distribution of mean κ values derived from 10,000 permutations.

Nearly all individual κ values (except for one participant in the CSR vs RW comparison for the degree metric) fall below zero, indicating within-subject disruption of the connectivity graph across sessions. This disruption is characterized by a reduced role of nodes that previously served as hub connector nodes, and an increased role of non-hub regions. For illustrative purposes, the disruption of the connectivity graph in two representative participants from our cohort is shown in Figure 6. These example connectograms highlight condition-related variability in functional network organization across different sleep states, but also interindividual differences in connectivity patterns. The latter provided motivation for adapting the HDI to a within-subject framework, allowing for more sensitive detection of subject-specific network disruption.

**Figure 6.**
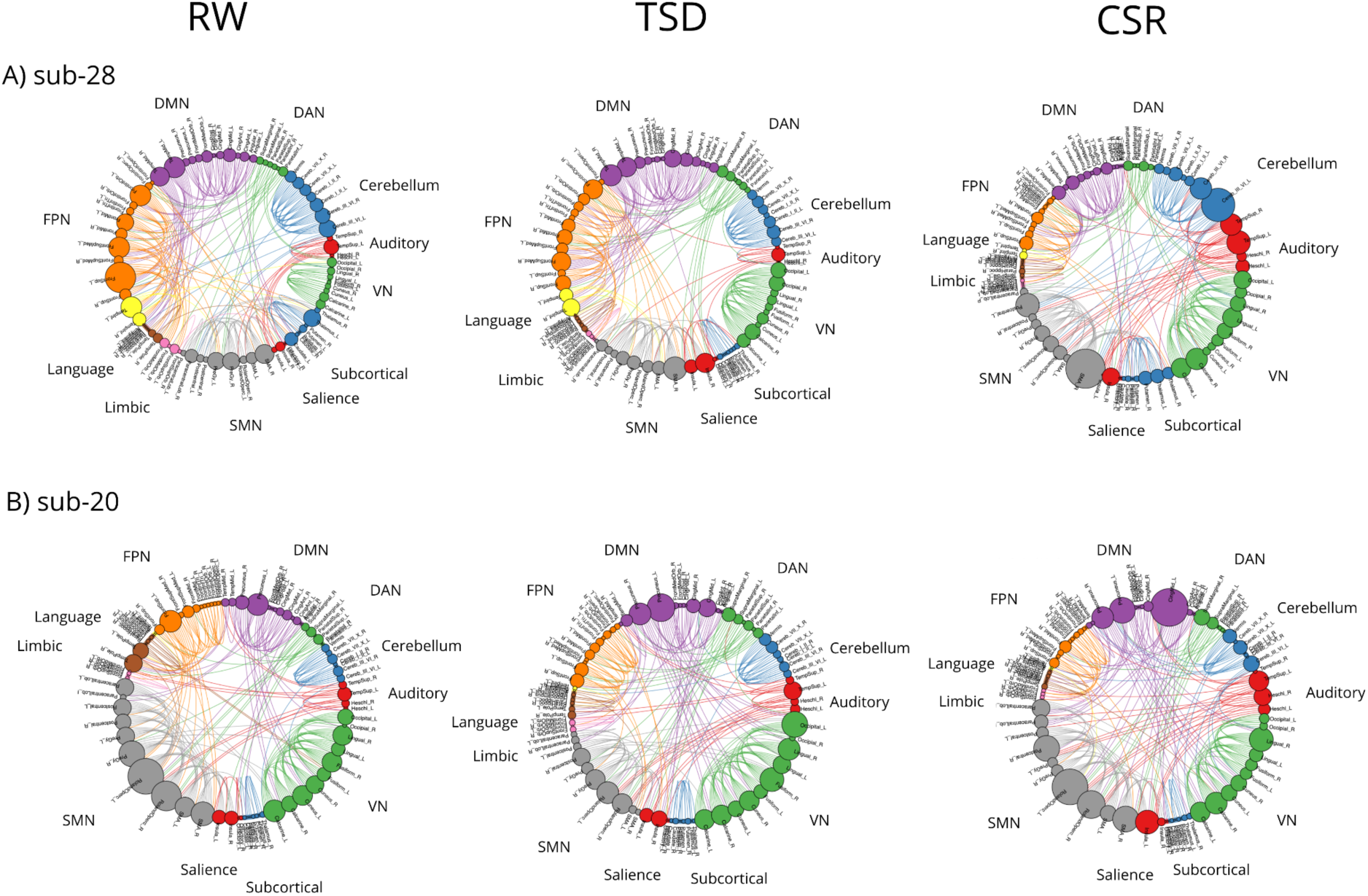
The connectograms of two example participants from our cohort in three sleep conditions. These chord diagrams show the functional connections (200 connections) within and across the sensorimotor network, visual network, frontoparietal network, default mode network, dorsal attention network, subcortical, auditory, language, limbic and cerebellum networks. Each of 89 brain areas from modified AAL atlas is represented by a dot with size corresponding to the number of connections of this node. Color coding is applied to enhance the visibility of specific networks. Abbreviations: RW - Rested Wakefulness, TSD - Total Sleep Deprivation, CSR - Chronic Sleep Restriction, DMN - Default Mode Network, DAN - Dorsal Attention Network, VN - Visual Network, SMN - sensorimotor network, FPN - frontoparietal network.

Charts for all of the participants are presented in the Supplementary Material S9.

#### Covariate-Constrained Manifold Learning (CCML)

The CCML revealed a spatial gradient along the first embedding dimension. Using degree centrality as input nodal metric and κ values as covariates, participants after TSD were predominantly located on the left side of the manifold, corresponding to more negative κ values, whereas the same participants after CSR tended to cluster toward the right. Importantly, this gradient was not imposed by group labels but instead, emerged from the covariate-driven structure of the data, with HDI κ values (derived from degree centrality) shaping the learned manifold (Figure 7).

**Figure 7.**
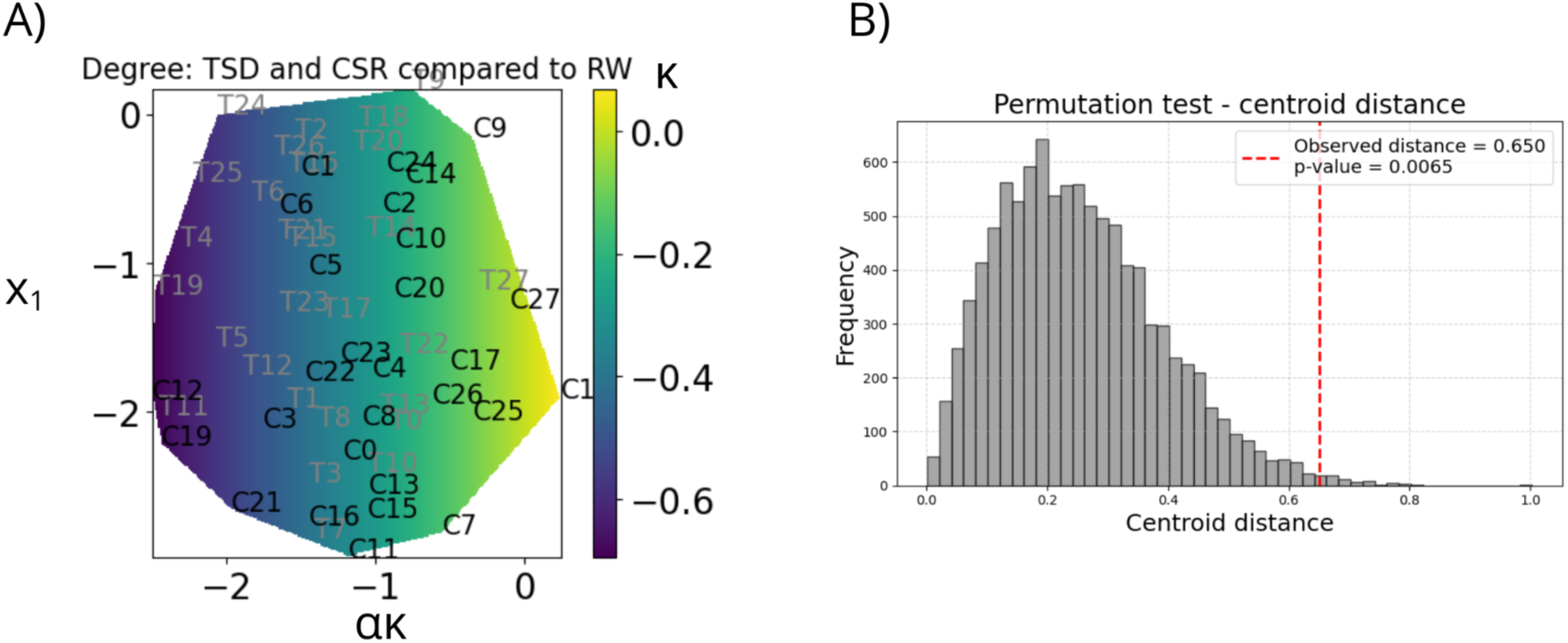
Covariate mapping onto the reduced space given by CCML, using degree centrality as a graph metric, and κ value from Hub Disruption Index analysis as covariate. A) Each point represents one participant, labeled as either TSD (T) or CSR (C). Covariate value (κ) is color coded. The coordinates correspond to [*α*κ; *xi*]^T^, where κ is the constrained variable and *xi* is the free parameter. A spatial gradient is visible, with TSD participants clustering toward the left (more negative κ) and CSR participants toward the right (less negative κ). While clusters show partial overlap, a clear tendency toward separation is observed. B) Permutation test of the observed Euclidean distance between group centroids in the reduced space. The distribution represents 10,000 centroid distances computed after random shuffling of group labels. The red dashed line indicates the real observed centroid distance (0.650), which lies in the far tail of the null distribution, yielding a permutation p-value of 0.0065. This confirms that the observed separation is unlikely to have arisen by chance.

Although the separation is not perfect, the two populations form visibly distinct, though partially overlapping, clusters. To statistically evaluate the separation between TSD and CSR conditions in the CCML reduced space, we computed the Euclidean distance between their group centroids and compared this observed value to a null distribution generated via 10,000 label permutations. The observed centroid distance was 0.650, significantly greater than expected by chance (permutation *p* = 0.0065), indicating that the two conditions exhibit a reliable, albeit not perfect, separation in the low-dimensional embedding. These CCML results are a confirmation of what we observed using within-subject HDI in TSD vs CSR comparison.

CCML was not performed using the clustering coefficient as input due to the presence of zero values in several nodes. Results of the CCML analysis based on closeness centrality are shown in Supplementary Materials S10.

#### Subjective sleepiness

We found that higher subjective sleepiness (KSS) is related to: lower global efficiency (*β* = -0.0045, *p* = 0.022), greater average path length (*β* = 0.0673, *p* = 0.027), greater graph distance between communities (*β* = 0.1381, *p* = 0.025) in RW condition. Collectively, these effects indicate reduced functional integration with increasing sleepiness. There were no significant relationships between both TSD and CSR, and subjective sleepiness of participants in these conditions. Detailed results are shown in Supplementary Table S13.

#### Sleep-related and circadian traits

In the RW condition, sleep quality (PSQI) was negatively associated with brain network modularity (*β* = -0.0356, *p* = 0.00089). This indicates that general poorer subjective sleep quality was linked to less modular (i.e., less segregated) brain networks in the RW condition. Moreover, across sessions, AM was positively related (*β* = 0.095, *p* = 0.0094) to the κ values obtained based on clustering coefficient in TSD vs RW comparison, indicating that individuals with higher distinctness of the circadian rhythm exhibit smaller disruptions after acute TSD. Moreover, we observed similar pattern in CSR vs RW comparison - higher amplitude was related to less negative κ values from clustering coefficient (*β* = 0.065, *p* = 0.057), however this should be considered a trend-level finding. Furthermore, we found that ME was negatively related to the HDI κ value for CSR vs TSD comparison based on degree (*β* = -0.064, *p* = 0.0211) and closeness (*β* = -0.059, *p* = 0.0466), indicating greater differences between CSR and TSD conditions among more evening-oriented individuals. Detailed results are shown in Supplementary Tables S14 and S15.

## Discussion

In this study, we aimed to investigate the effects of sleep deprivation on functional connectivity in the brain using fMRI and graph-theoretical methods. Each participant underwent scanning across three sessions, corresponding to three sleep conditions: rested wakefulness, acute total sleep deprivation, and chronic sleep restriction. To our knowledge, this is the first studies to investigate both acute TSD and CSR within the same experimental framework, and the very first to enable a direct comparison between these two sleep loss conditions using a within-subject, longitudinal design.

We found that both TSD and CSR are associated with a reorganization of the topological structure of brain functional networks (Figure 5). We observed consistently negative κ values, indicating that brain regions serving as hubs during RW, lost their hub status after sleep deprivation, while less central nodes gained connections. Moreover, all of the κ values in the direct comparison between CSR and TSD were also negative, which strongly suggests that these two forms of sleep loss lead to the different network disruptions, and therefore affect brain network organization possibly through distinct underlying mechanisms. This hypothesis is further supported by the results from CCML method, which revealed a separation between the CSR and TSD conditions when using dimensionality reduction, with the HDI-derived κ values included as covariates (Figure 7).

Importantly, all of these observed effects were reliable regardless of the graph cost used (5% - 200 edges, 10% - 400 edges, 15% - 600 edges), the atlas applied (modified anatomical AAL with 89 regions or modified functional AICHA with 391 regions), or the nodal metric analyzed (degree centrality, closeness centrality, and clustering coefficient). This robustness provides strong evidence for a systematic reorganization of brain functional networks not only under conditions of TSD and CSR relative to RW, but also between the two sleep deprivation conditions themselves.

In addition, we identified differences between conditions in nodal metrics using linear mixed-effects models. Several brain regions exhibited significant changes, however, we note that the specific nodes identified varied slightly depending on the graph cost used. Nevertheless, the overall pattern remained consistent. For clarity and comparability, we present detailed results based on the 10% graph cost in the main text (Figure 4) and discuss these findings below - while acknowledging that this method is inherently sensitive to the thresholding applied during the graph construction process. Results obtained using different costs are provided in the supplementary materials S11.

Previous research consistently demonstrated that acute TSD disrupts the brain’s intrinsic functional connectivity, particularly within the Default Mode Network (DMN). In the well-rested state, the DMN typically shows anticorrelated activity with task-positive networks. Following TSD, studies reported reduced connectivity inside DMN and between DMN and its related anticorrelated network [19]. Regions of the DMN (such as the medial prefrontal and posterior cingulate cortex) become less synchronized with each other, indicating the DMN cannot maintain its normal integrity under conditions of sleep deprivation. In our study we found node-level connectivity changes that correspond to these observations. We report reduced connectivity in Left Angular Gyrus and the bilateral Posterior Cingulate Cortex (PCC) but also increased connectivity in the Right Superior Parietal Lobule and the bilateral Precuneus. This apparent dissociation suggests that different subsystems of the DMN may be differentially affected by acute TSD. Indeed, recent work supports this interpretation. Chen et al. identified dorsal and ventral subnetworks within the DMN and observed a dissociable effect: acute TSD was related to the lower connectivity in the dorsal DMN (which includes midline posterior hubs like the PCC and angular gyrus) but increased functional connectivity within the ventral DMN [64]. This aligns very well with results described here. Our observation of dorsal DMN disruption (reduced connectivity in PCC and angular gyrus) alongside increased degree in the precuneus and superior parietal lobule (functionally closer to ventral DMN) fits this dissociation model well. Other studies corroborate this phenomenon. For example, Gujar and colleagues reported that a single night of sleep deprivation disrupts the typical task-related deactivation pattern of the DMN, resulting in a double dissociation between anterior and posterior midline regions of this network [65]. Taken together, these findings support the hypothesis that some DMN nodes become hypoactive while others may remain active or even hyper-connected to compensate for the negative effect of acute sleep loss. Notably, Chen et al. further demonstrated that lower functional connectivity in dorsal DMN was associated with impairments in basic cognitive performance, whereas individuals who exhibited increased connectivity between the subsystems of DMN, showed better preserved working memory. This suggests that such an enhancement may serve a compensatory mechanism in the brain’s response to sleep deprivation [64].

Regarding CSR, we found reduced connectivity in the Left Angular Gyrus and Right Inferior Parietal Lobule when compared to the RW state. These two regions are considered as part of the DMN, and salience network, respectively. Due to novel approach, i.e. within-subject, longitudinal design, we were able to directly compare between two distinct types of sleep loss. Interestingly, we found that CSR when compared to TSD is associated with decreased connectivity in the Frontoparietal network (FPN) - in the Left Inferior Frontal Gyrus (triangular part). This may suggest that this region is particularly vulnerable to CSR, while it remains relatively preserved or less disrupted following TSD. We know from prior studies that the prefrontal-driven networks are among the most affected by sleep loss in general [66,67]. Although several studies have shown decreased prefrontal activity following TSD [68,69], others have reported preserved or even increased activation [70,71], likely reflecting task-related compensatory mechanisms [15]. These inconsistencies in the literature highlight the importance of considering both types of sleep loss when interpreting connectivity changes in the prefrontal cortex.

Based on our results, cerebellum also plays a crucial role in the brain’s response to the sleep loss state. Following TSD, we observed reduced degree in the sensorimotor region of the left cerebellum (lobules III–VI) and an increased clustering coefficient in the right cerebellar lobules III–VI in comparison to RW, indicating region-specific shifts in local connectivity. In the comparison between CSR and TSD, we found higher connectivity in the cerebellum after CSR, specifically in bilateral lobules III–VI (sensorimotor regions) and in the left lobules VII–X, which are associated with cognitive and limbic functions [72]. These findings are supported by the study of Zhang et al., who reported altered cerebellar functional connectivity after 36 hours of TSD [73], further highlighting the sensitivity of cerebellar networks to acute sleep loss, but not to the chronic one.

From a global network perspective, Ben Simon et al. showed profound reduction in network modularity following acute TSD, evident in the limbic, default-mode, salience and executive modules [74]. On the other hand, CSR also degrades overall connectivity efficiency. Farahani and colleagues showed that global topology of brain networks changed after a week of sleep at ∼35% below normal sleep need - the characteristic path length increased and “small-world” index decreased relative to the well-rested state [14]. All these findings indicate a loss of functional segregation within the brain and a shift towards a more random-like network [19,21]. While we did not observe a significant reduction in modularity following TSD or CSR compared to the RW condition, we did find a significant decrease in modularity accompanied by a trend-level association with reduced average clustering and average graph distance after CSR when compared directly to the TSD state. This suggests qualitative differences in how the two forms of sleep loss alter global brain network organization.

Taken together, all these observations derived from within-subject HDI, CCML, nodal and global metrics, support the hypothesis that acute TSD and CSR affect brain networks through distinct mechanisms, with the cerebellum possibly serving as a key structure in differentiating these neural responses.

Additional contribution of this study is the adaptation of the classical HDI method to suit our longitudinal design. Specifically, instead of using group-level control averages as the reference (as in the original HDI formulation), we used each participant’s own control scan as an individual reference. As illustrated in Figure 6 and Supplementary Figure S9, we observed variability in functional connectivity not only across different sleep conditions, but also substantial interindividual variability even within the RW baseline state. This observation motivated us to move away from a group-average control, allowing for a more precise within-subject comparison of functional network changes across sleep states. This modifications appears to be more sensitive to subject-specific disruptions of functional connectivity as reflected in the consistency of our results. We observed uniformly negative κ values across all comparisons, graph costs, and atlases. This indicated a robust and replicable pattern of brain network reorganization and, together with the permutation tests we conducted (Figure S5), supports the validity of our methodological modification of HDI.

Finally, it is important to address the methodological choices involved in graph construction, including the graph cost and the brain atlas used for parcellation. In this study, we primarily present results obtained using modified AAL atlas and graphs with 400 edges, corresponding to a 10% cost threshold - a commonly adopted approach in functional connectivity analyses [46,75]. However, it is worth noting that there is currently no golden standard regarding optimal graph density in brain network research. Similarly, there is no consensus on the choice of brain atlas for parcellation. To confirm the robustness of our results and to rule out the possibility that they were specific to a particular graph cost or brain atlas, we conducted supplementary analyses using alternative graph costs (5% and 15%, corresponding to thresholds of 200 and 600 edges, respectively) as well as an additional atlas - a modified version of the AICHA atlas, which is functionally rather than anatomically defined.The results of these complementary analyses are provided in the Appendix S11, S12. For several nodes, we observed similar significant differences in graph metrics across multiple thresholds (e.g., Cerebellum, Precuneus, Angular Gyrus), which may indicate robust and consistent disruptions in network properties under sleep deprivation conditions. These threshold-independent effects strengthen the interpretation that the observed changes are not merely artifacts of specific graph density choices but reflect meaningful alterations in brain connectivity. However, the fact that some different nodes emerge at different thresholds, reflects the changes in the graph’s structure [46]. As the number of edges increases, the network tends toward a lattice-like structure where node properties converge and spurious edges may be included. The HDI is the only method that consistently identifies differences at the whole-brain level, highlighting the distributed nature of sleep deprivation’s impact. This suggests that while regional node metrics capture local changes, they may miss the systemic reorganization of the entire network.

Using the modified AICHA atlas with 391 regions yielded nodal-level results consistent with those from AAL only for the degree metric, and brain regions: Cerebellum and Cingulate Cortex. This discrepancy may be due to several factors, including: (I) AICHA’s being functional atlas rather than anatomical (as AAL atlas), (II) higher number of smaller regions, which are more sensitive to noise [38], and (III) higher number of comparisons inherent to finer parcellation requires more stringent multiple comparison correction, which may obscure potentially meaningful changes in nodal metrics. Although both atlases describe the same brain system, increasing the number of regions alters the graph’s structure, influencing topological properties and statistical power. Still, the fact that HDI produced analogous results with both atlases supports the idea of a global network disruption following sleep loss, different mechanisms behind TSD and CSR, and further confirm the stability and robustness of the within-subject HDI.

In addition to objective graph-theoretical measures of neural responses to sleep loss, we evaluated subjective measures, classified as state-level and trait-level. Subjective sleepiness was assessed in each experimental condition and treated as a state-level measure. In the RW condition, greater sleepiness was associated with general reduced functional integration - lower global efficiency, longer path length, and greater between-community graph distance. In other words, self-reported sleepiness predicted objective features of the brain’s functional network and is associated not only with subjective experience but also with measurable differences in brain activity. These associations were not observed after TSD or CSR, likely because those experimental conditions uniformly elevated sleepiness, reducing between-participant variance. Among trait-level features, poorer sleep quality was associated with lower modularity, meaning the brain’s functional network was less clearly partitioned into specialized communities. Subjective amplitude of the circadian rhythm was positively related to κ values derived from the clustering coefficient for TSD vs RW. A similar, trend-level association was present for CSR vs RW comparison, indicating that individuals with a more pronounced circadian rhythm, stronger preserve brain’s local network structure after both sleep loss types. Although extreme chronotypes were excluded from our cohort, higher ME scores were related to more negative κ values in CSR vs TSD comparisons (based on both, degree and closeness), indicating that more evening-oriented individuals experience greater differences between these two types of sleep loss conditions. These patterns align with prior reports showing that chronotype modulates vulnerability to sleep loss [76,77]

Overall, our results support the hypothesis that distinct mechanisms underlie the changes in brain functional connectivity after TSD and CSR. However, these findings can also be interpreted in the context of the work by Van Dongen and colleagues (2003), who showed that multiple nights of partial sleep restriction can accumulate, effectively impairing cognition almost as severely as total sleep loss, especially for attention and working memory [22]. In our study, CSR involved five consecutive nights of restricted sleep, therefore, future research might extend the length of the CSR period to investigate the hypothesis of cumulative effect.

Moreover, Farahani and colleagues showed that the effects of CSR vary depending on the time of day and its interaction with individual circadian rhythms [14,78]. Time of the day, chronotype and subjective amplitude of circadian rhythm (distinctness), are known to influence brain activity and structure [61,62,79–81]. In our study, we attempted to control for these factors by scheduling all scanning sessions during the same daytime window (between 9:30 a.m. and 3:30 p.m.) and by excluding participants with extreme morning or evening chronotypes. Nonetheless, we found significant associations between circadian rhythm characteristics and global graph metrics. A natural continuation of our study would be to enlarge the cohort to include individuals with extreme chronotypes and a wider range of circadian rhythm amplitudes, to further examine factors that may modulate functional brain connectivity and neural responses to different types of sleep loss.

Finally, in our CCML analysis, we focused on distinguishing between TSD and CSR conditions, however, future studies could explore linking the second axis of the embedding to specific cognitive or psychological features.

## Limitations

The primary limitation of the present study is the relatively small sample size. Although the inclusion of 28 participants is consistent with sample sizes commonly reported in fMRI research, it nonetheless limits the statistical power and generalizability of the findings. A key strength of the study, however, lies in its longitudinal design: each participant underwent three scanning sessions, resulting in a total of 84 fMRI scans. This within-subject approach enhances the reliability of the observed effects. Nevertheless, replication with larger samples is necessary to substantiate these results.

A further limitation concerns the restricted age range of the sample (20-36 years). This range was deliberately selected to minimize the influence of age-related variability in brain function. While this approach enhances internal validity, it constrains the generalizability of the findings to other age groups. Future studies should aim to include participants across a broader age spectrum to evaluate the developmental or aging-related dynamics of the observed neural processes related to sleep deprivation.

## Conclusions

This study reveals that acute total sleep deprivation and chronic sleep restriction distinctly reorganize brain functional networks, as shown by graph-theoretical fMRI analysis in a within-subject, longitudinal design. Both sleep-deprived states were associated with negative κ values derived from the within-subject Hub Disruption Index, indicating a loss of centrality in previous network hubs. Crucially, direct comparisons between TSD and CSR uncovered condition-specific network disruption. This divergence was further supported by a significant separation in Covariate-Constrained Manifold Learning embeddings, suggesting these two types of sleep loss may affect brain network topology through different mechanisms. These findings were robust across multiple graph costs (thresholds), atlases, and nodal metrics, highlighting their reliability. By adapting the HDI to use an individual’s own scan as reference, we accounted for interindividual variability and confirmed consistent, condition-specific network changes. Significant changes in degree centrality, closeness centrality, and clustering coefficient were observed in regions of the default mode network, frontoparietal network, and cerebellum. Finally, subjective sleepiness was associated with reduced network integration, poorer sleep quality with lower modularity, and circadian phenotype marked individual sensitivity to sleep loss. Together, these findings provide strong evidence that TSD and CSR induce qualitatively different alterations in functional brain organization, with the cerebellum emerging as a potential differentiating structure.

## Code and data availability

Time series, atlases and code used in this study are available in the following GitHub repository: https://github.com/PatrycjaScislewska/sleep_deprivation_graphs/

All preprocessing steps can be fully reproduced using the following GitHub repository: https://github.com/veronicamunoz/rs_graph_processing

## Acknowledgements

We gratefully acknowledge all study participants for their dedication and compliance with the study procedures, particularly for their consistency in adhering to the sleep deprivation protocol.

## Disclosure statement

### Preprint repositories

Manuscript appeared as a preprint in the BioRxiv repository.

## Authors’ contributions

All authors conceived the project. All authors interpreted the results and revised the manuscript. P.S.: Methodology, Formal analysis, Code development, Writing - Original Draft, Visualization; A.C.V.: Methodology, Formal analysis, Writing - Original Draft; I.S.: Writing - review & editing, Supervision; H.K.O.: Conceptualization, Methodology, Formal analysis, Study design, Data collection, Funding acquisition; S.A.: Methodology, Formal analysis, Code development, Writing - review & editing, Supervision; A.D.: Conceptualization, Study design, Data collection, Methodology, Funding acquisition, Writing - review & editing, Supervision

## Supplementary materials

S1. Modified version of AAL atlas description. Text reproduced from [39], Supplementary Materials:

Modified version of classical AAL parcellation scheme The classical AAL parcellation scheme is composed by 116 regions including the cerebellum. We have merged some of the regions, reducing the parcellation to 89 regions. Merged regions are: frontal medial orbital and rectus (one region for left and one for right hemisphere); occipital superior, middle and inferior (one region for left and one for right hemisphere); temporal pole superior and medial (one region for left and one for right hemisphere); the cerebral crus (one region for left and one for right hemisphere); areas III, IV, V and VI of cerebellum (one region for left and one for right hemisphere); areas VII, VIII, IX, X of cerebellum (one region for left and one for right hemisphere) and finally, the vermis (one single region for both hemispheres).

S2. Quality check of the fMRI data after motion regression (following the approach of Power and colleagues [41]). A) For each frame of data in one subject, the framewise displacement (FD) of a frame of data is plotted against the absolute values of the differentials of RS fMRI timecourses of 89 ROIs. A locally weighted regression (LOESS) curve (black line) was fit to the relationship between FD and BOLD signal change using a span parameter of 0.02, enabling the visualization of fluctuations and potential motion-related artifacts. B, C, D) Identically produced LOESS curves from all 28 subjects in all three scanning sessions are plotted against FD. These results confirm that there is no significant relationship between motion and the BOLD signal in the described dataset.

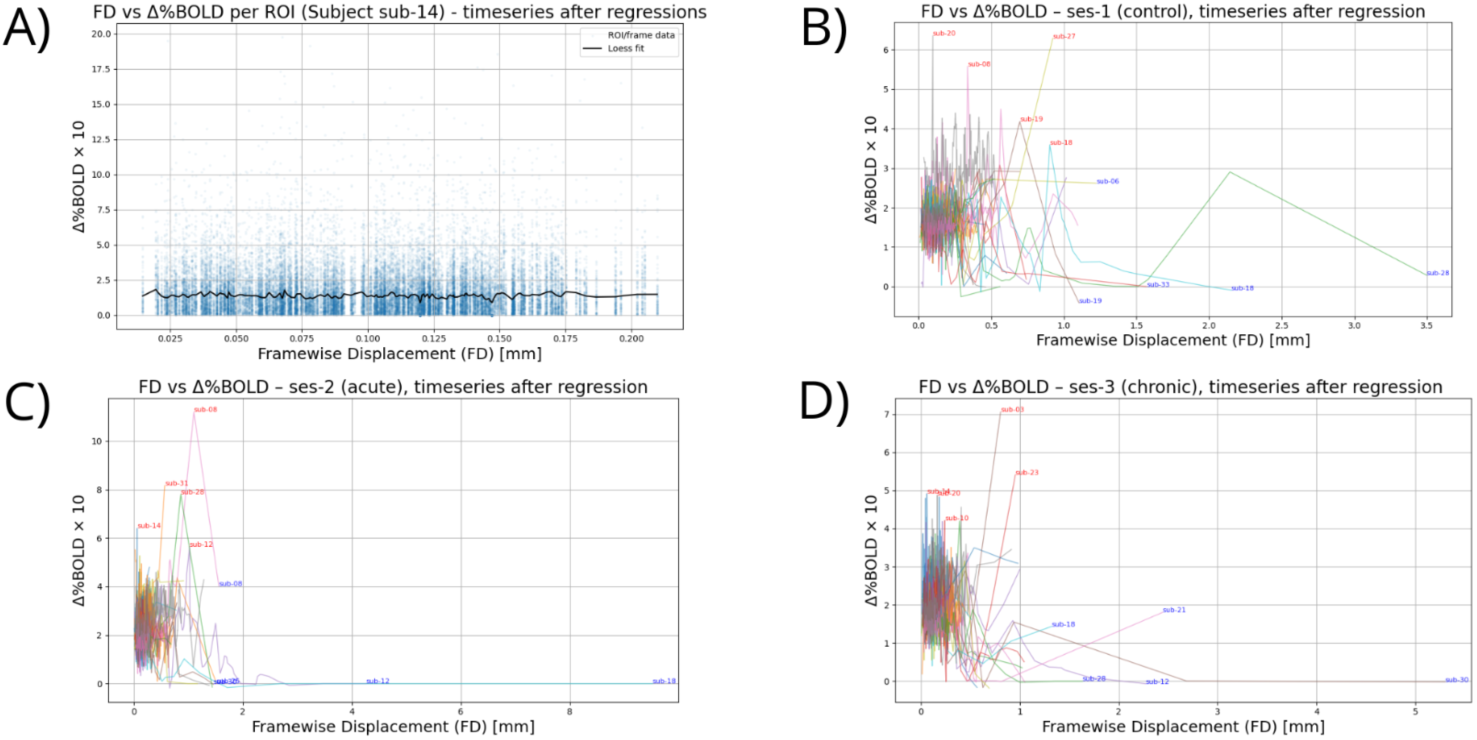

S3. Boxplots with numbers of significant edges per participant per scanning session for both atlases (modified AAL and AICHA). Dots represent participants. Each plot is annotated by the subject’s ID with the smallest number of significant edges.

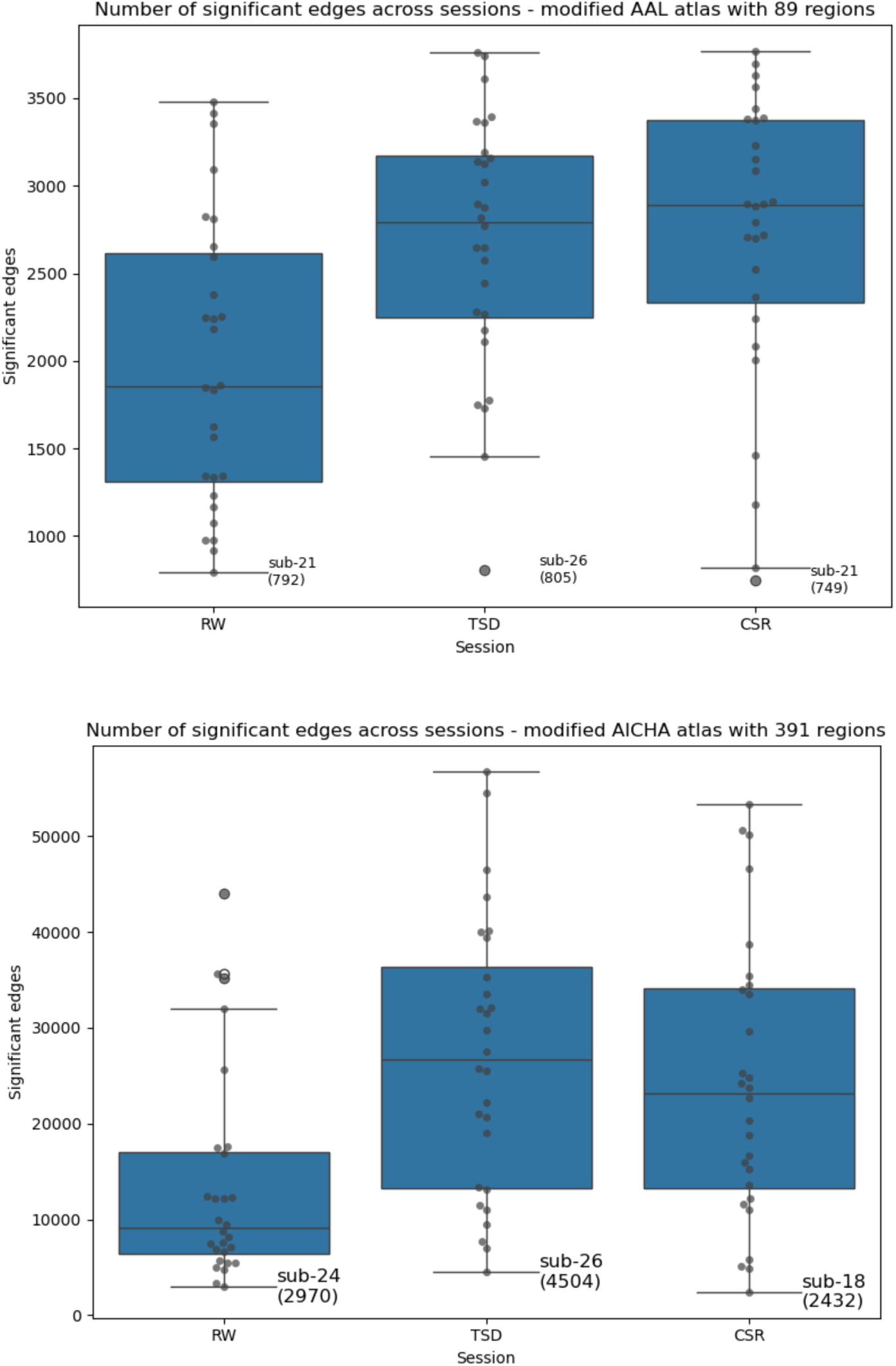

S4. Equations for nodal metrics:

The simplest graph metric is the degree of a node *i*, D*_i_*. It corresponds to the number of edges connected to a node *i*

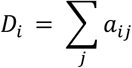

where 𝑎_*ij*_, is the entry in the adjacency matrix *A*, with 𝑎_*ij*_ = 1, if nodes *i* and *j* are connected, and 0 otherwise. Degree centrality is normalized by dividing degree value by the highest node degree in the network.

We also tested the clustering coefficient Cc_i_, which quantifies how nodes in a graph tend to cluster together and form local groups [52]. The clustering coefficient is defined as the ratio between twice the number of connections among neighbors of node *i* and the total number of possible connections between those neighbors.

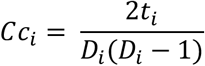

where *t_i_* is the number of connections among neighbors of node *i*.

The third considered metric is closeness centrality, which reflects how quickly the information can be spread through the network, and is calculated as the inverse of the average shortest distance from the node to all other nodes in the graph. [51].

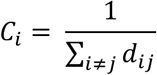

where d_ij_ is the shortest path distance between nodes *i* and *j*.

S5. Schematic representation of permutation tests performed to evaluate the within-subject HDI method. A) Original data: participants scanned under different conditions (here: reference condition (marked in orange) and condition *t* (marked in blue)) with correct within-subject pairing and condition labels. Dots represents the nodes, color code (light red, vivid red, dark red) represents example degree values. B) Node permutations: within-subject pairing and condition labels are preserved, but degree of nodes (here: colors of nodes) is randomly shuffled in the condition *t* scans. C) Unpaired permutations: correct condition labels, but participants’ order in the condition *t* scans is randomly mixed across all subjects, breaking within-subject pairing. D) Paired permutations: condition labels are randomly assigned, preserving the pairing but not the original condition assignments.

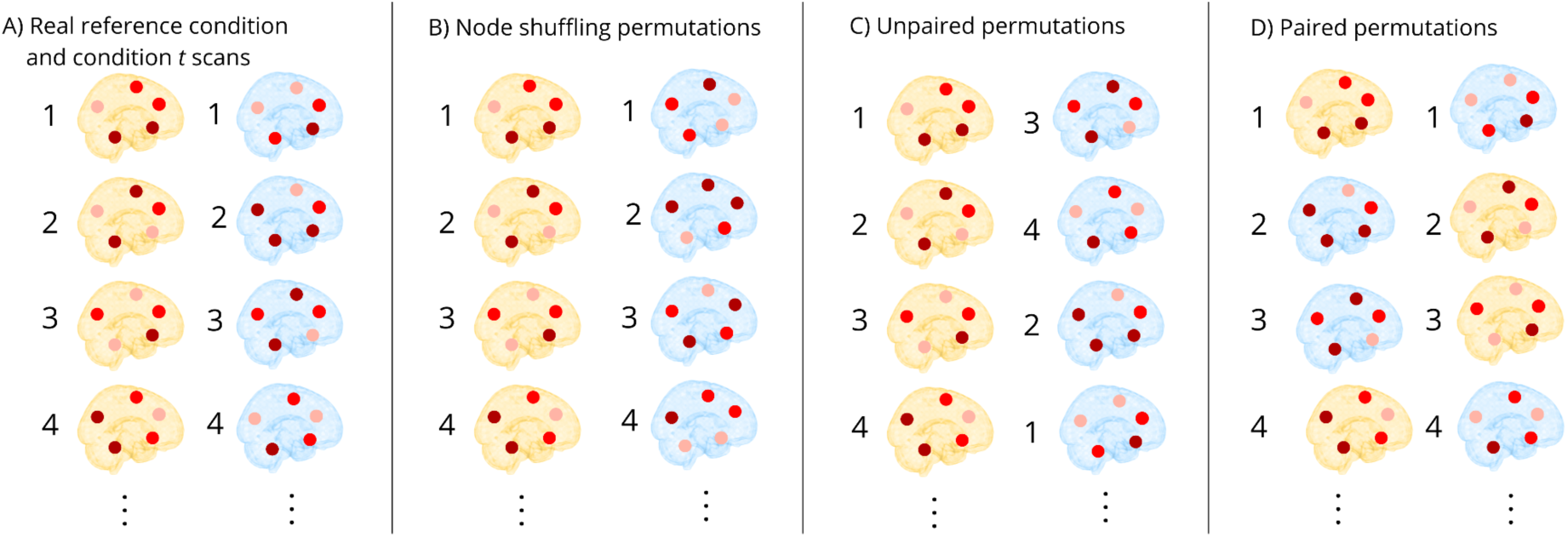

S6. Results of the Node shuffling permutations test and Unpaired permutations test.

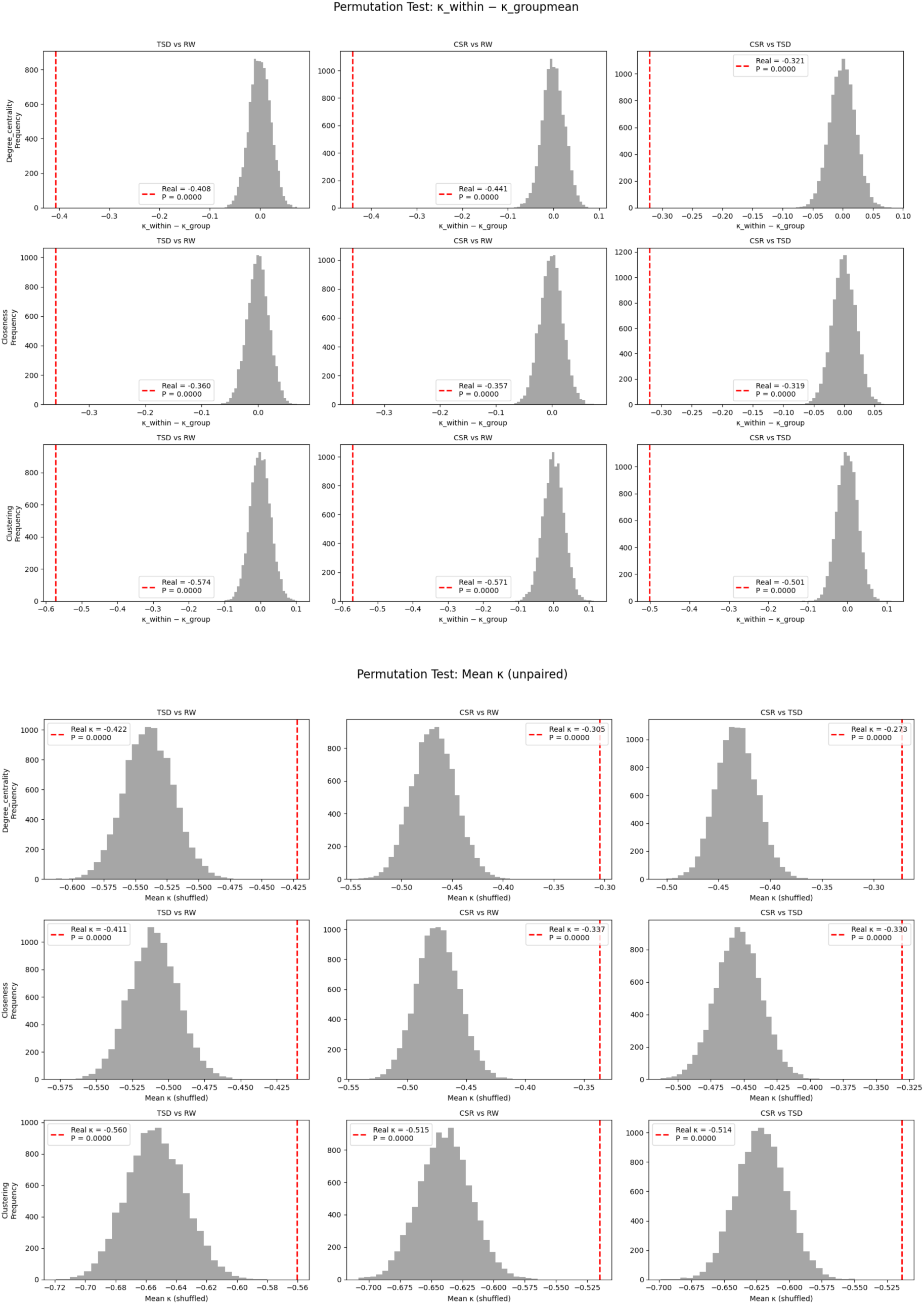

Results of Paired permutations test are provided in the main text, Figure 5.

S7. Table with details from linear mixed-effects model for global metrics. Bolded and underlined text indicates significant results (p < 0.05), bolded text indicates trend-level associations.

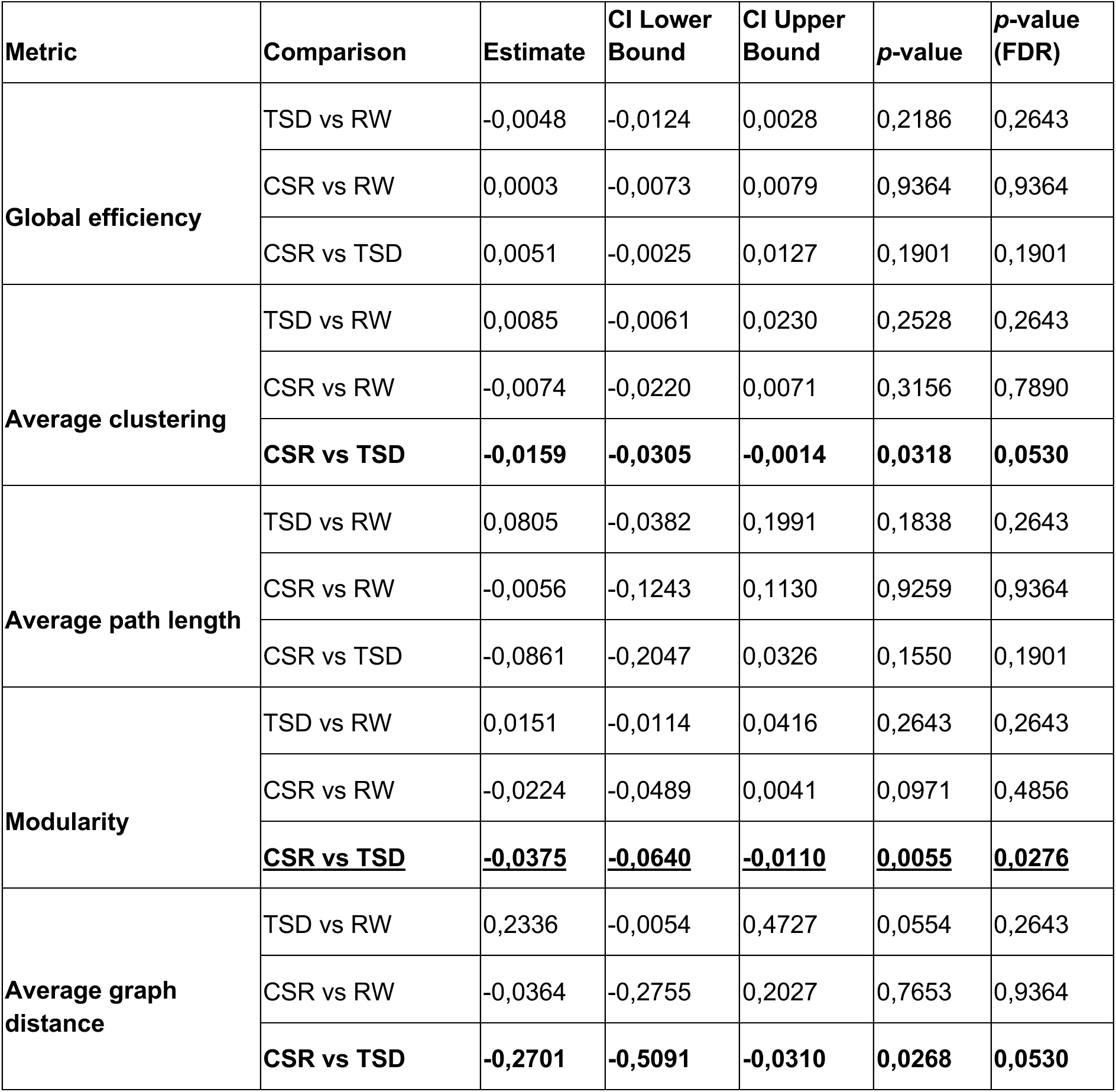

S8. Permutation tests to check the robustness of results of the linear mixed-effects model analysis for global metrics.

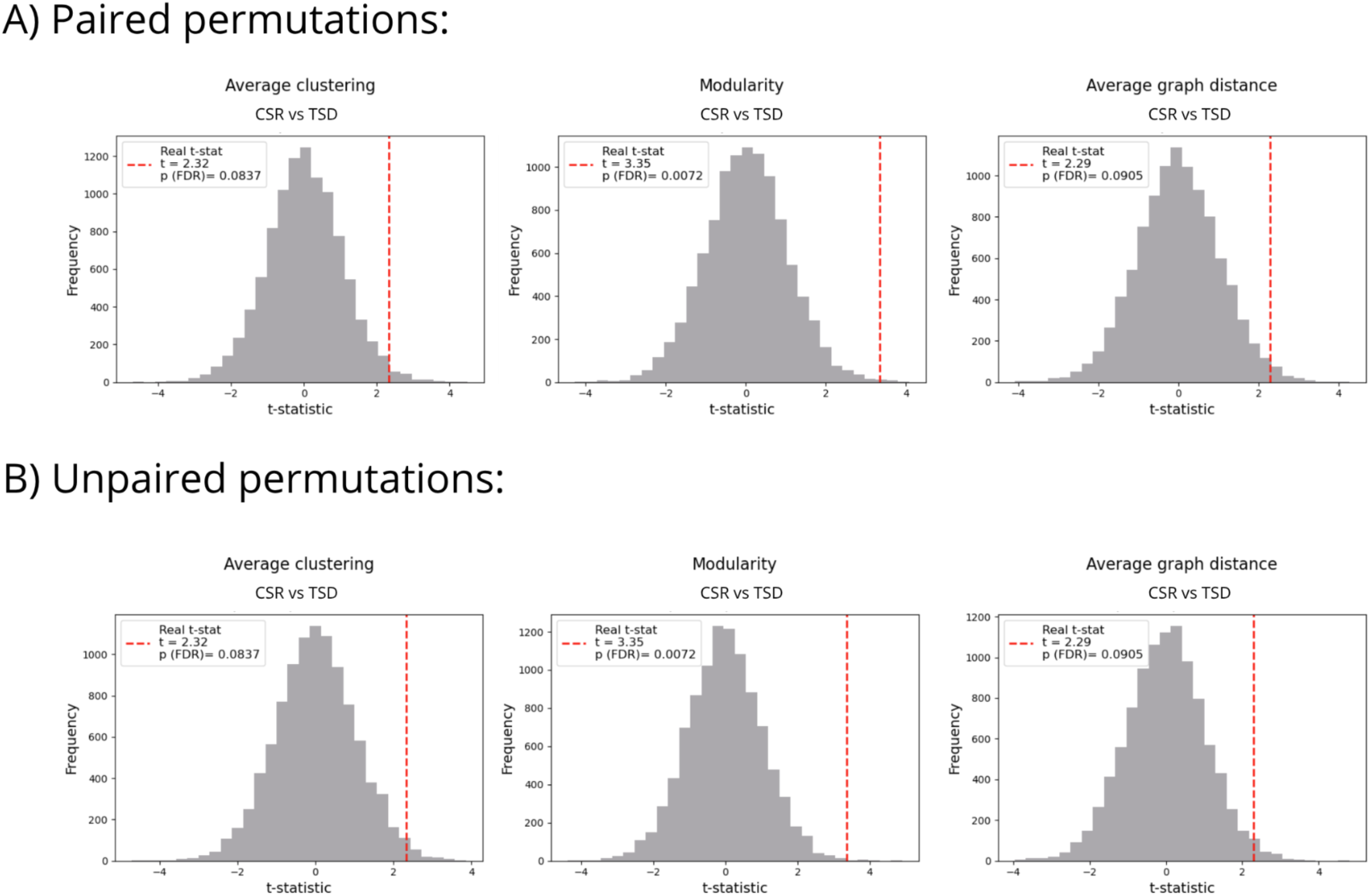

S9. Connectograms of the 28 participants in our cohort.

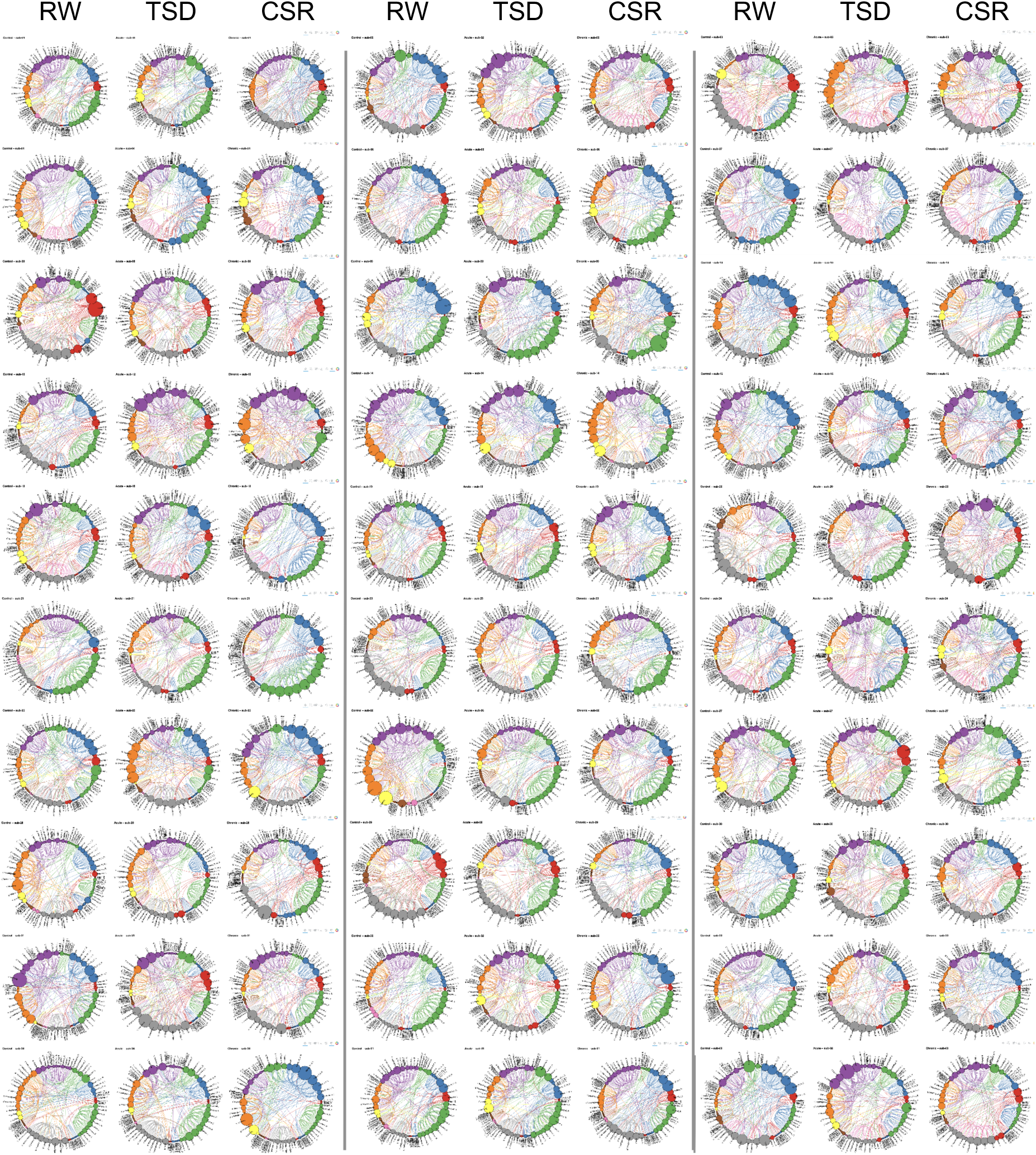

S10. Results of the CCML analysis based on closeness centrality

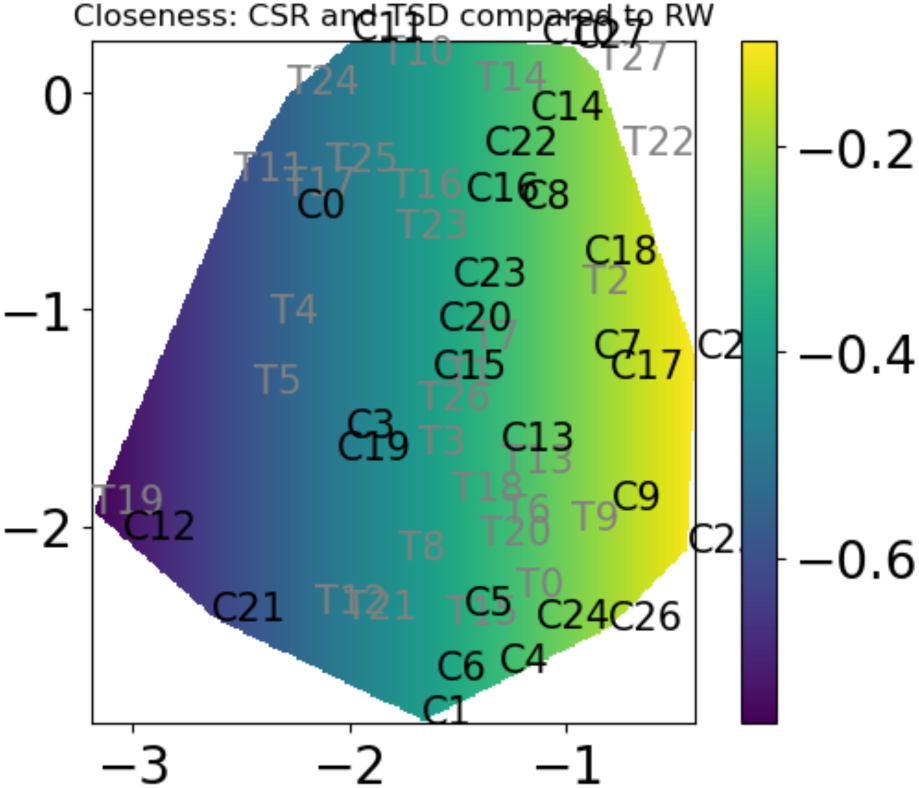

S11. Results for additional graph densities. Graphs were constructed at multiple edge densities by thresholding the correlation matrices to retain a fixed number of strongest connections. Specifically, we tested 15 edge thresholds ranging from 200 to 3000 edges (in steps of 200).

To confirm the robustness of our result, here we report results obtained using graphs constructed with 200 and with 600 edges. All statistical analyses were performed identically to those described in the main text of this publication.

Graph with 200 edges (5% cost):

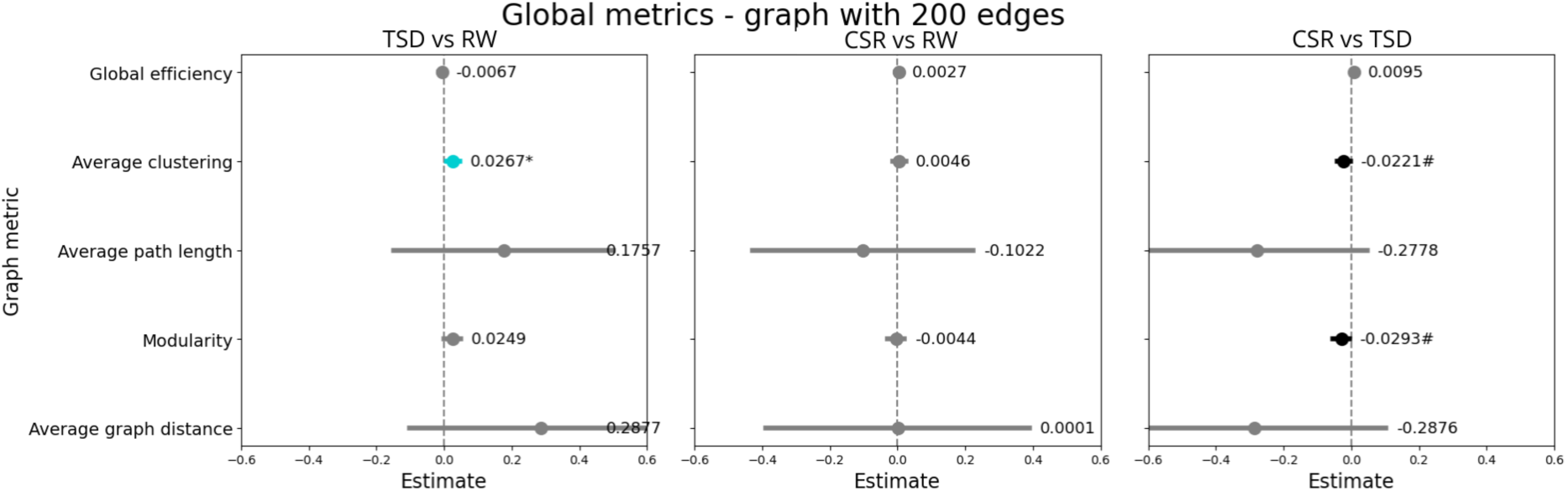

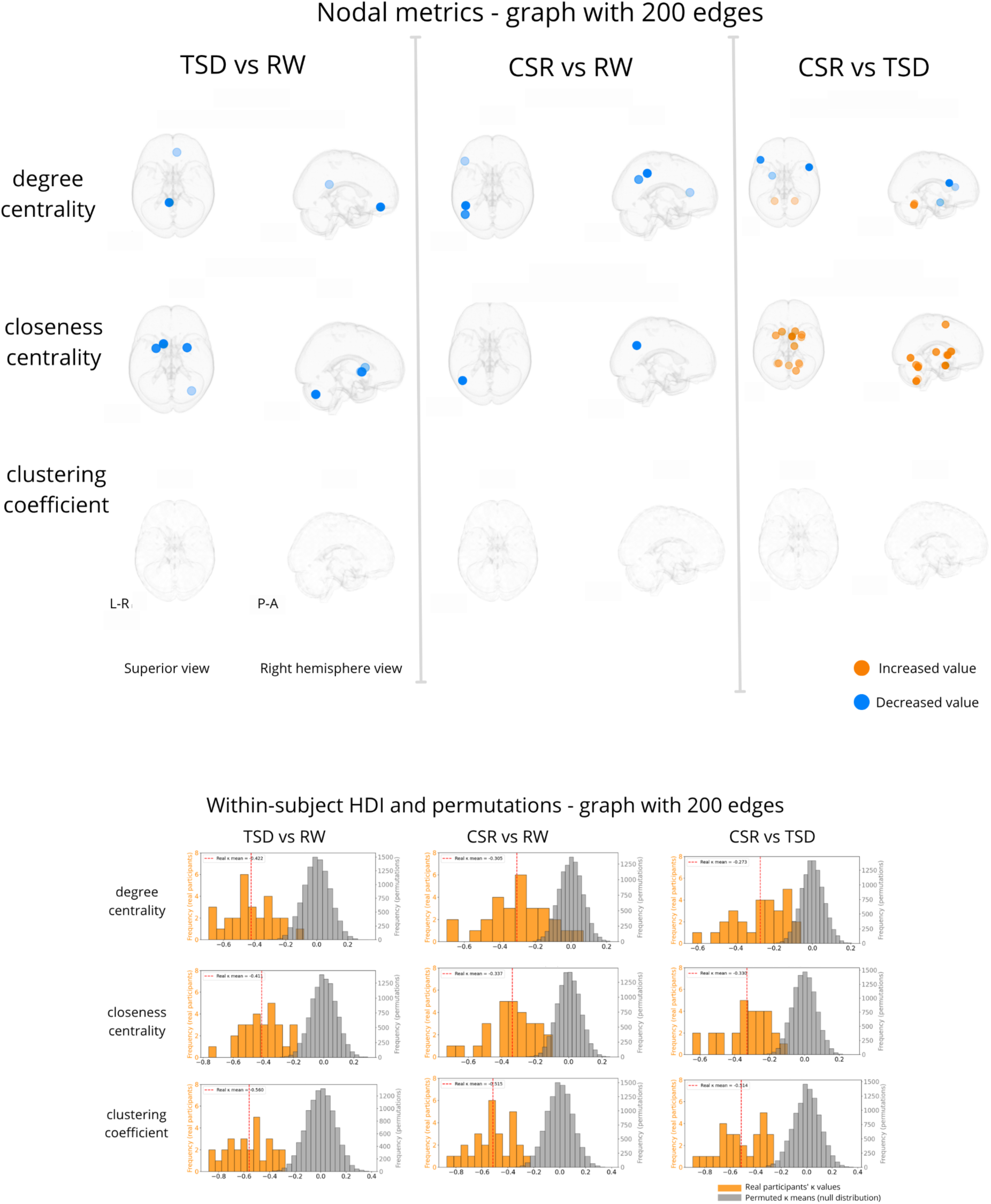

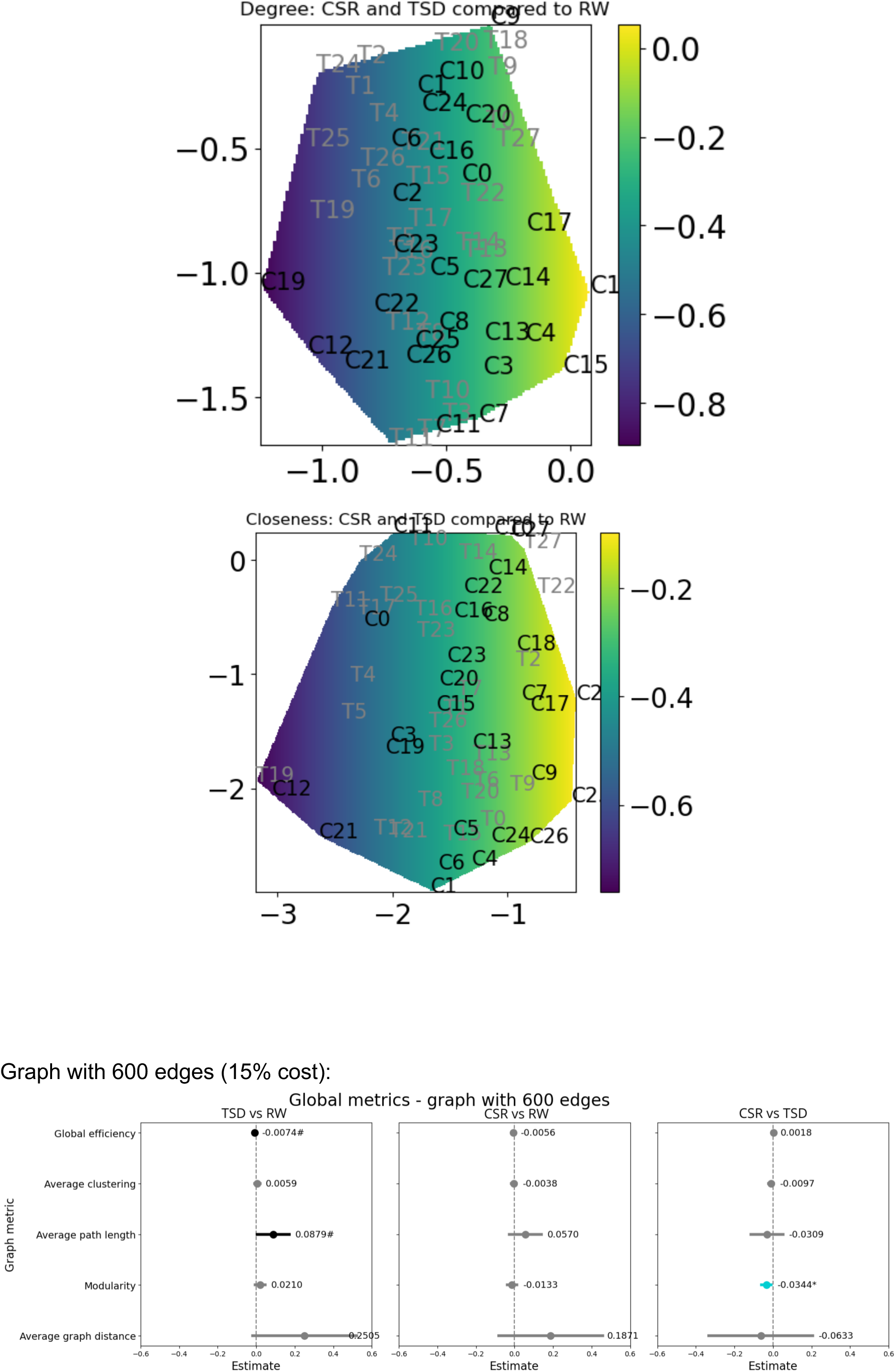

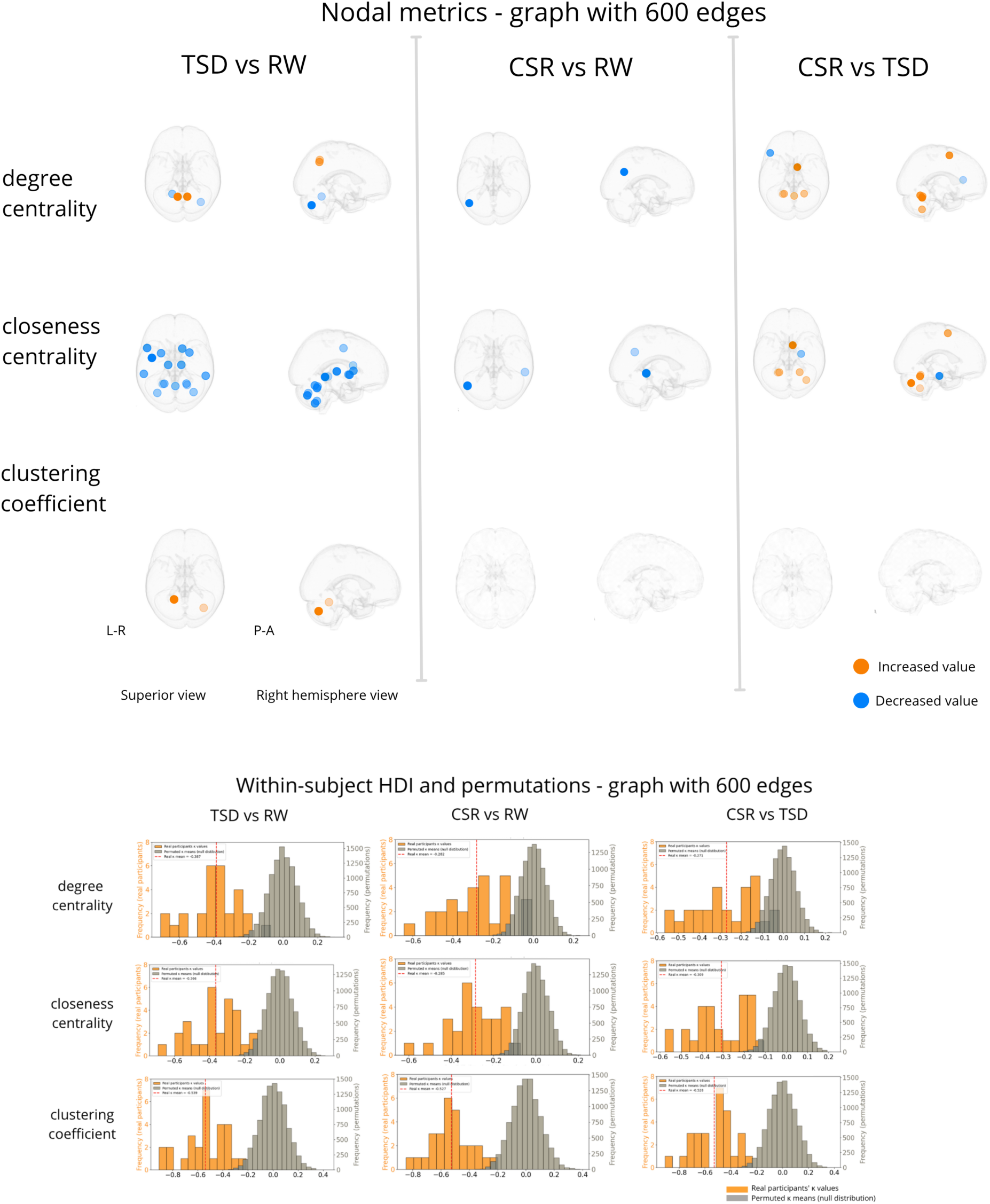

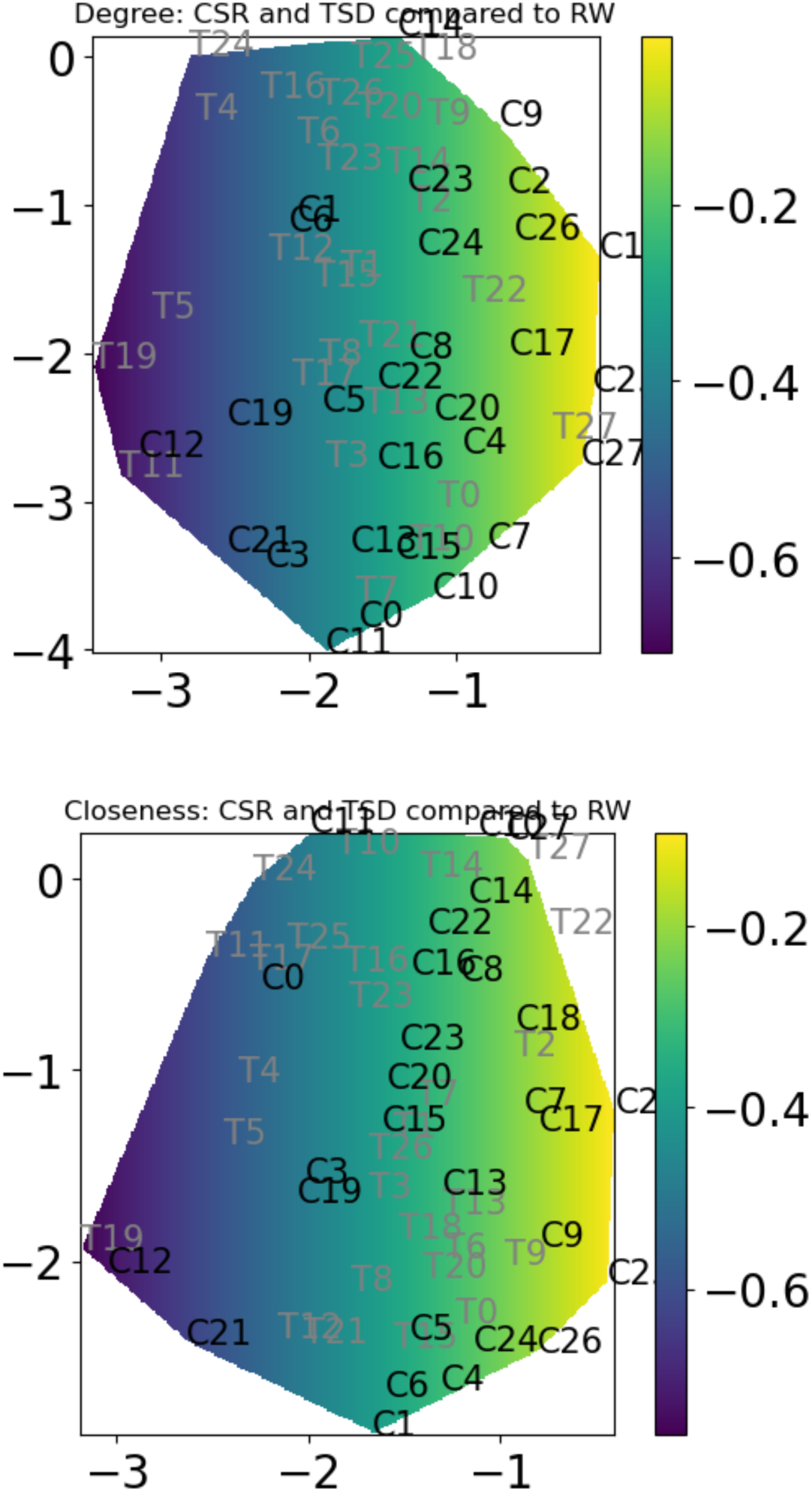

S12. Results for modified AICHA atlas (added cerebellum from AAL atlas) with 391 regions in total:

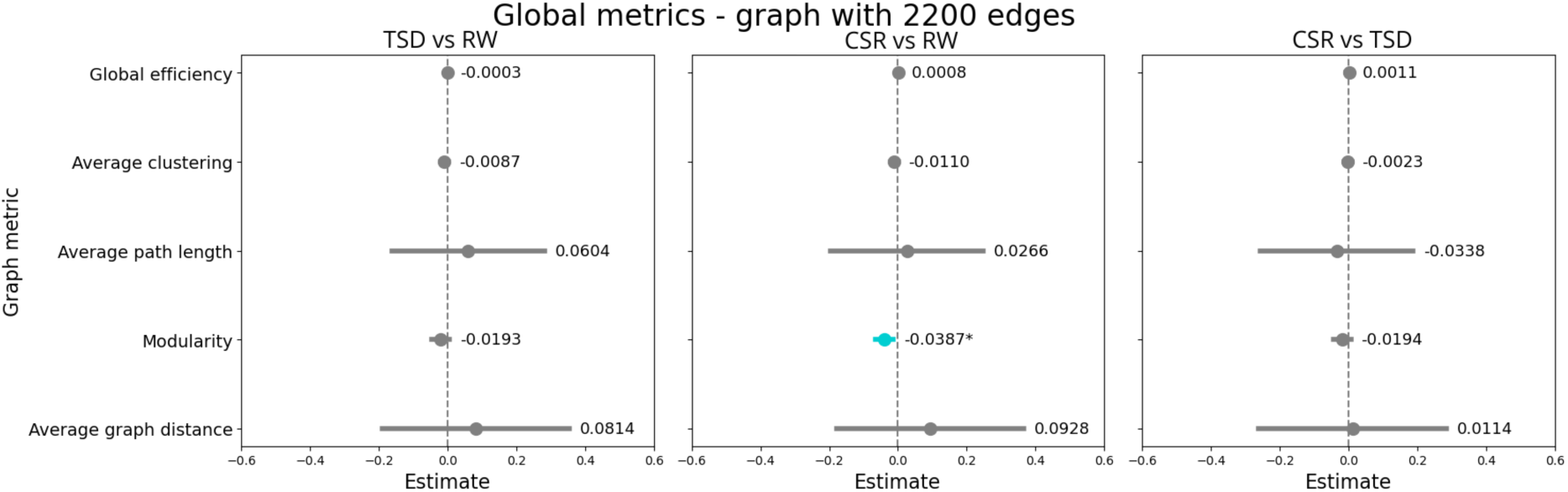

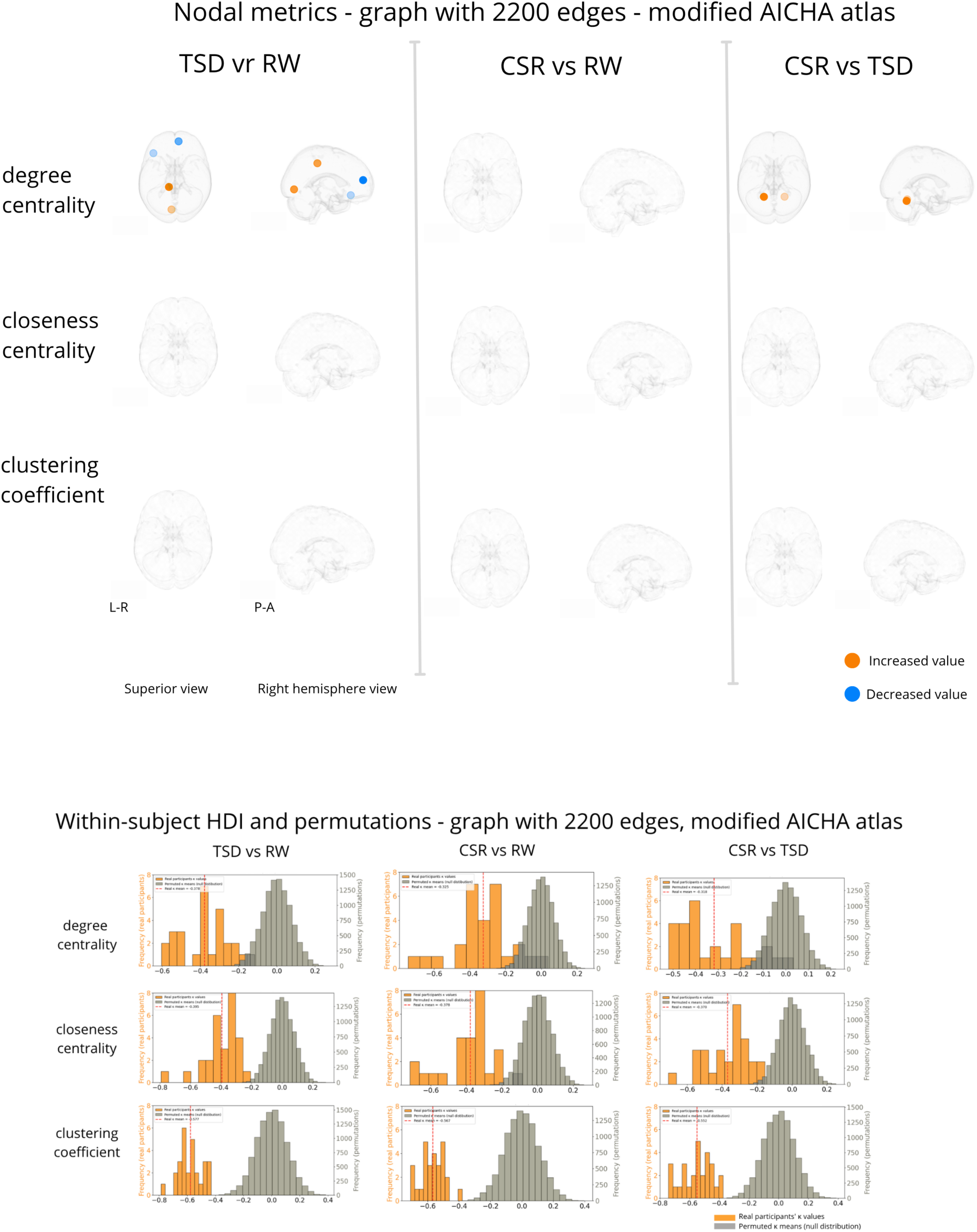

S13. Results of the linear Mixed-Effects Model. Our goal was to verify if the subjective sleepiness is associated with global graph metrics (global efficiency, average clustering coefficient, average shortest path length, modularity, and average graph distance) in each of our experimental conditions (RW, TSD, CSR).

| Global metric | Condition | $\beta$ | z | CI_low | CI_high | p-value FDR |
| --- | --- | --- | --- | --- | --- | --- |
| Global efficiency | **<u>RW</u>** | **<u>-0.0045</u>** | **<u>-2.6765</u>** | **<u>-0.0078</u>** | **<u>-0.0012</u>** | **<u>0.0223</u>** |
|  | TSD | -0.0011 | -0.7797 | -0.0039 | 0.0017 | 0.5189 |
|  | CSR | 0.0011 | 0.6451 | -0.0022 | 0.0044 | 0.5189 |
| Average clustering | RW | 0.0001 | 0.0206 | -0.0066 | 0.0067 | 0.9836 |
|  | TSD | -0.0037 | -1.3247 | -0.0093 | 0.0018 | 0.4383 |
|  | CSR | -0.0031 | -1.0533 | -0.0088 | 0.0027 | 0.4383 |
| Average shortest path length | **<u>RW</u>** | **<u>0.0673</u>** | **<u>02.6097</u>** | **<u>0.0168</u>** | **<u>0.1179</u>** | **<u>0.0272</u>** |
|  | TSD | 0.0242 | 01.938 | -0.0192 | 0.0675 | 0.4111 |
|  | CSR | -0.0049 | -0.1874 | -0.0565 | 0.0467 | 0.8514 |
| Modularity | RW | 0.0031 | 0.4902 | -0.0092 | 0.0154 | 0.8494 |
|  | TSD | -0.0047 | -0.8930 | -0.0150 | 0.0056 | 0.8494 |
|  | CSR | -0.0013 | -0.1899 | -0.0142 | 0.0117 | 0.8494 |
| Average graph distance | **<u>RW</u>** | **<u>0.1381</u>** | **<u>02.6410</u>** | **<u>0.0356</u>** | **<u>0.2405</u>** | **<u>0.0248</u>** |
|  | TSD | 0.0250 | 0.5969 | -0.0572 | 0.1073 | 0.5880 |
|  | CSR | 0.0290 | 0.5417 | -0.0758 | 0.1337 | 0.5880 |

S14. Results of the OLS model. Our goal was to verify if trait-level sleep and circadian measures predict brain global graph metrics.

| Global metric | Predictor | $\beta$ | R<sup>2</sup> | p-value FDR |
| --- | --- | --- | --- | --- |
| average clustering | AM | -0.0074 | 0.1665 | 0.5367 |
|  | ME | 0.0035 | 0.1665 | 0.6699 |
|  | PSQI | -0.0150 | 0.1665 | 0.2101 |
| average path length | AM | -0.0115 | 0.0135 | 0.8354 |
|  | ME | -0.0083 | 0.0135 | 0.8354 |
|  | PSQI | -0.0287 | 0.0135 | 0.8354 |
| average graph distance | AM | -0.1272 | 0.0664 | 0.6718 |
|  | ME | -0.0347 | 0.0664 | 0.6979 |
|  | PSQI | -0.0781 | 0.0664 | 0.6718 |
| global efficiency | AM | 0.0010 | 0.0258 | 0.7964 |
|  | ME | 0.0011 | 0.0258 | 0.7964 |
|  | PSQI | 0.0024 | 0.0258 | 0.7964 |
| modularity | AM | -0.0181 | 0.3456 | 0.0744 |
|  | ME | 0.0099 | 0.3456 | 0.2304 |
|  | **<u>PSQI</u>** | **<u>-0.0356</u>** | **<u>0.3456</u>** | **<u>0.0009</u>** |

S15. Results of the OLS model. Our goal was to verify if trait-level sleep and circadian measures predict differences in the functional connectivity between our experimental conditions.

| Nodal metric | HDI comparison | Predictor | $\beta$ | $R^2$ | p-value_FDR |
| --- | --- | --- | --- | --- | --- |
| Degree centrality | $\kappa_{TSD\_vs\_RW}$ | AM | 0.0189 | 0.0858 | 0.7897 |
|  |  | ME | -0.0434 | 0.0858 | 0.7897 |
|  |  | PSQI | -0.0112 | 0.0858 | 0.7897 |
| | $\kappa_{CSR\_vs\_RW}$ | AM | 0.0064 | 0.1519 | 0.8280 |
|  |  | ME | -0.0345 | 0.1519 | 0.5546 |
|  |  | PSQI | -0.0677 | 0.1519 | 0.1136 |
| | $\kappa_{CSR\_vs\_TSD}$ | AM | 0.0207 | 0.1894 | 0.6677 |
|  |  | **ME** | **-0.0635** | **0.1894** | **0.0211** |
|  |  | PSQI | -0.0149 | 0.1894 | 0.7089 |
| Closeness centrality | $\kappa_{TSD\_vs\_RW}$ | AM | 0.0016 | 0.0729 | 0.9589 |
|  |  | ME | -0.0063 | 0.0729 | 0.9589 |
|  |  | PSQI | -0.0369 | 0.0729 | 0.8683 |
| | $\kappa_{CSR\_vs\_RW}$ | AM | 0.0280 | 0.1878 | 0.4848 |
|  |  | ME | -0.0141 | 0.1878 | 0.6155 |
|  |  | PSQI | -0.0519 | 0.1878 | 0.2463 |
| | $\kappa_{CSR\_vs\_TSD}$ | AM | 0.0404 | 0.2345 | 0.2251 |
|  |  | **ME** | **-0.0594** | **0.2345** | **0.0466** |
|  |  | PSQI | -0.0089 | 0.2345 | 0.8083 |
| Clustering Coefficient | $\kappa_{TSD\_vs\_RW}$ | **AM** | **0.0946** | **0.3490** | **0.0094** |
|  |  | ME | 0.0076 | 0.3490 | 0.8902 |
|  |  | PSQI | 0.0049 | 0.3490 | 0.8902 |
| | $\kappa_{CSR\_vs\_RW}$ | **AM** | **0.0652** | **0.2096** | **0.0570** |
|  |  | ME | 0.0153 | 0.2096 | 0.9224 |
|  |  | PSQI | -0.0020 | 0.2096 | 0.9459 |
| | $\kappa_{CSR\_vs\_TSD}$ | AM | 0.0047 | 0.0084 | 0.9486 |
|  |  | ME | 0.0146 | 0.0084 | 0.9486 |
|  |  | PSQI | 0.0031 | 0.0084 | 0.9486 |

